# Ground tilt representation in the rodent cerebral cortex

**DOI:** 10.1101/2025.09.04.674164

**Authors:** Fumiya Imamura, Hiroto Imamura, Reiko Hira, Yoshikazu Isomura, Riichiro Hira

## Abstract

Information about ground tilt at one’s location is indispensable for integrating the body with its surrounding environment. However, how ground-tilt information, together with movement and posture information, is represented in the cerebral cortex remains unclear. To address this issue, we developed a new six-degree-of-freedom platform that can rapidly and precisely impart ground motion to head-fixed mice. Using the Diesel2p large field-of-view two-photon mesoscope, we performed calcium imaging to simultaneously capture activity dynamics across broad cortical areas from the frontal to the parietal cortex while driving the platform through various motion patterns. We found a core area for the ground-tilt information at the parietal cortex, although tilt-direction neurons were found across the dorsal cortex. Neuronal populations preserved tilt-direction information within a common low-dimensional manifold across three distinct motion conditions, indicating a condition-invariant tilt code. Furthermore, to investigate shared dynamics across multiple cortical areas, we introduced broadcast- subspace analysis as an extension of communication subspace analysis. This extended framework revealed that tilt information occupied the second principal shared axis, following movement information. These results reveal the core cortical area for processing tilt information, its universal format, and its sharing across the dorsal cortex, thereby providing a framework for cortical information processing that integrates movement, posture, and environment.

## Introduction

Animals in natural environments continually encounter ground tilt, and the rapid and precise estimation of its direction and magnitude is critical for stabilizing current posture and planning near-future movements. Integration of ground-tilt information involves not only the mechanical properties of the musculoskeletal system and reflex mechanisms in the brainstem and spinal cord, but also the cerebellum, basal ganglia, superior colliculus, and cerebral cortex (Takakusaki 2017; Deliagina et al. 2007; Jacobs and Horak 2007). Among these, cortical involvement in cross-hierarchical integration is particularly pivotal (Chiou et al. 2018; Adkin et al. 2006; Beloozerova et al. 2003; Drew, Prentice, and Schepens 2004; Boebinger et al. 2024; Solis-Escalante et al. 2019). Neurons that encode gravity direction and ground tilt based on vestibular input have been identified in parietal regions including primary somatosensory cortex (S1) and posterior parietal cortex (PPC) (Angelaki et al. 2020; Dougherty et al. 2021; Disse et al. 2023; Nandakumar et al. 2021). In rodents, the frontoparietal cortex including secondary motor cortex (M2) and PPC represents posture (Mimica et al. 2018; Soma et al. 2018), and postural and movement information is broadly shared between motor and sensory cortices (Mimica et al. 2023; Nandakumar et al. 2021). In monkeys, posture-dependent activity related to voluntary movement has been observed in several PPC regions (Lacquaniti et al. 1995; Buneo and Andersen 2006). In addition, during anticipatory postural adjustments, higher motor areas (e.g., M2) and the corticoreticular pathway from the frontal cortex to the reticular formation play central roles, enabling anticipatory responses to ground tilt through control of proximal musculature (Takakusaki 2017; Jacobs and Horak 2007; Adkin et al. 2006; Chiou et al. 2018; Boebinger et al. 2024; Solis-Escalante et al. 2019). These observations strongly suggest that ground-tilt signals are computed and shared within the parietal and frontal cortex and serve as foundational signals for postural and movement control. Despite their importance, it remains unknown whether ground tilt is represented independently of posture and movement within the cortex, how it is functionally localized, and to what extent it is shared across cortical networks.

Recently, it has been reported that specific low-dimensional subspaces shared across population activity mediate communication between cortical areas (Semedo et al. 2019; MacDowell et al. 2025; Gonzalez et al. 2024). Activity related to body movement is widely observed throughout the cortex and likely pervades large-scale cortical dynamics (Stringer et al. 2019; Musall et al. 2019). Surveying eight parietal regions and analyzing the correlation structure of their population activity, we found that the largest shared subspace was common to all pairs of areas (Hira et al. 2024). Moreover, in recordings from the fronto-parietal cortex, the dominant shared component predominantly reflected movement-related signals (Imamura et al. 2025). At the same time, cortical activity can covary with fluctuations in motivation, salience, and arousal (Raut et al. 2023; Reitman et al. 2023; Stringer et al. 2019). Intriguingly, cortico-cortical communication between M2 and PPC includes working memory and temporal information (Voitov and Mrsic-Flogel 2022; Imamura et al. 2025), suggesting that subspaces are governed by factors beyond movement. We therefore hypothesize that, because information about ground tilt and posture is distributed across the cortex, these signals form dominant non-motor components of the cortex-wide shared dynamics. To test this hypothesis, it is essential to comprehensively examine wide cortical areas using large-scale, multi-area recordings while precisely manipulating ground tilt as an exogenous signal and simultaneously monitoring posture and movement.

For this purpose, we developed a new six-degree-of-freedom arbitrary ground-motion platform (hereafter referred to as **6AGM**). By combining this with the Diesel2p mesoscope (Hira et al. 2024; Yu et al. 2021; Hira et al. 2025), we mapped neurons that selectively encode the direction of ground tilt. As a result, direction-selective neurons were localized near the border between S1tr and S1hl, yet were also observed across all dorsal cortical areas. Intriguingly, these neuronal populations represented tilt direction on a circular manifold that remained invariant across different motion patterns. We further developed an analysis method, termed GCARP (generalized canonical component ablation-based representational probing), to extract information shared across all regions of the dorsal cortex. Applying this method revealed that the shared dynamics contained the strongest contribution from movement, followed by ground tilt and posture. Taken together, these findings indicate that the cerebral cortex contains not only regions specialized for ground-tilt processing, but also a distributed architecture in which ground tilt, posture, and movement information are broadly shared.

## Results

### Construction and evaluation of the platform

To examine posture and movement and their modulation by ground tilt, we built a fully open-source platform, 6AGM (**Fig. 1a**). The platform is assembled from custom-made components (**Fig. 1b; Supplementary Information**) and is controlled by four modules that enable immediate movement to arbitrary targets: Position-Sampling Module, Feedback Module, Calculation Module, and Voltage-Output module. Working sequentially, these modules drive the platform precisely to any specified pose (**Fig. 1c**). Motion is generated by six stepper motors located beneath the platform, each of which adjusts the position of a support column along its linear guide. The resulting platform pose is defined by six variables: the 3D center coordinates (X, Y, Z) and the rotation angles (pitch, roll, yaw), which are linked to the actuator positions through the platform’s kinematics 𝑓 **(Fig. 1d)**. To move the platform to a target pose, the control system solves the corresponding inverse kinematics 𝑓^−1^ in the Calculation Module, determining the required actuator positions.

**Figure 1.**
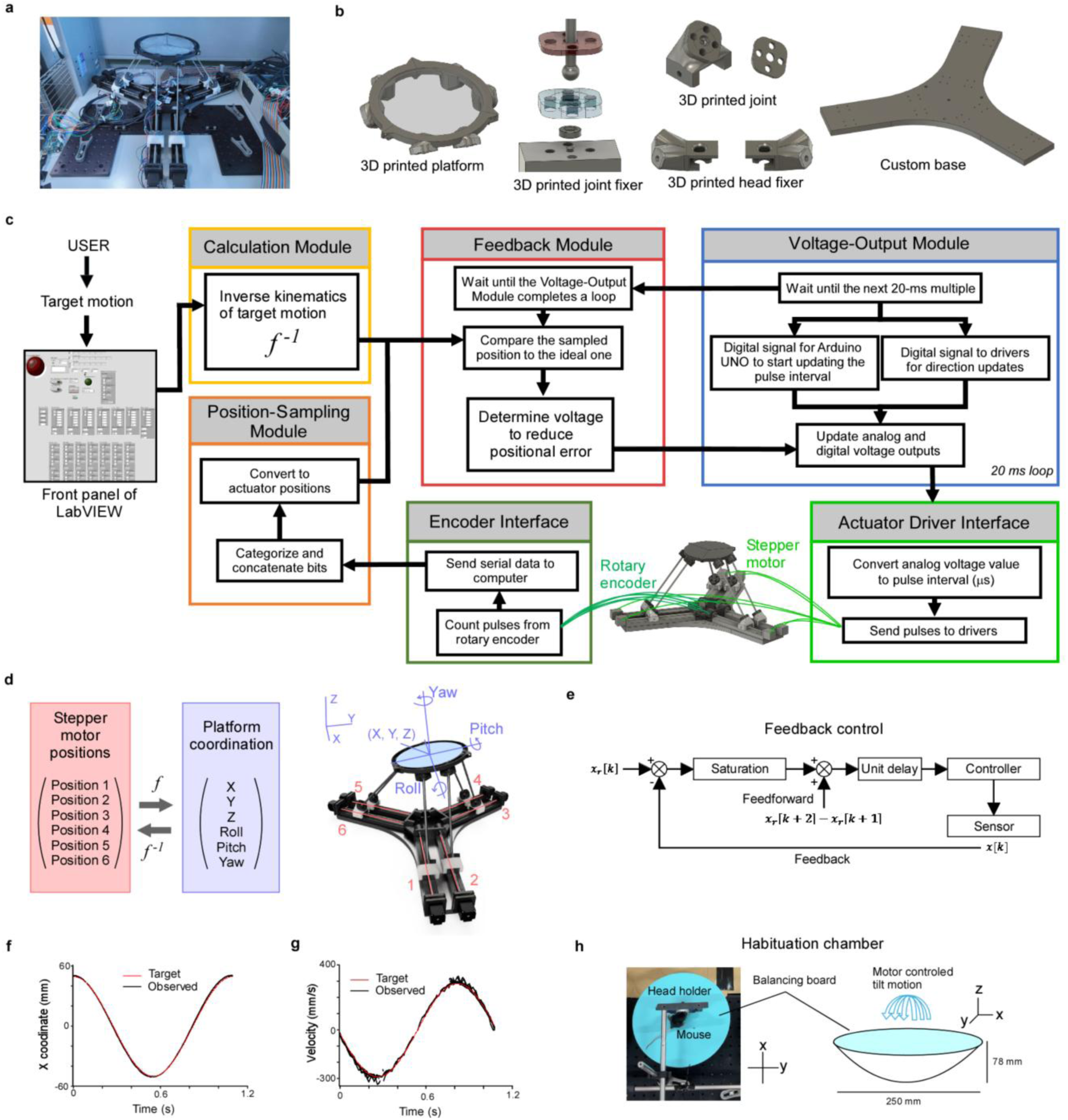
Development of a new 6-degree-of-freedom platform and evaluation of its accuracy. **a.** Photograph of the new 6-degree-of-freedom arbitrary ground-motion platform (6AGM). **b.** Components of 6AGM (for details, see **Supplementary Information**). **c.** Schematic of platform control. Four modules (Position-Sampling Module, Feedback Module, Calculation Module, Voltage-Output Module) move the platform rapidly and sequentially toward a target pose. The Position sampling module receives precise stepper-motor positions via the encoder interface, and the Voltage output module drives the stepper motors via the actuator driver interface. **d.** The six stepper motors control six pose parameters. The platform pose is defined by the 3D coordinates of the center (X, Y, Z) and the rotation angles (pitch, roll, yaw). A unique mapping 𝑓 relates actuator states to platform pose, and the inverse mapping 𝑓^−1^ is used for control, computed in the Calculation module. **e.** Schematic of the feedback control. Key features: (i) the current motor positions are read and the error relative to the target pose is computed at each update; (ii) the error is clamped to a maximum value to avoid overcompensation; (iii) the combined feedback and feedforward commands yield fast, accurate convergence with minimal overshoot and smooth trajectories. **f.** Target position (red) and observed platform position (black). **g.** Target velocity (red) and observed platform velocity (black). **h.** Habituation apparatus. To acclimate mice to ground motion, head-fixed animals were placed on an unstable balance board.

We implemented a feedback control mechanism to achieve smooth motion (**Fig. 1e**). The controller first reads the current stepper positions and computes the error relative to the current target. A feedforward control signal then predicts the change from the next step to the subsequent step. These two signals determine the next motor output. By assigning an upper bound (“saturation”) on the error to prevent instability, the platform follows accurate and fast trajectories for arbitrary tilts, rotations, and translations (**Fig. 1f,g**). The control software was written in LabVIEW and released as open-source software. To habituate mice to platform motion, we constructed an apparatus with an unstable board and performed habituation under head-fixation (**Fig. 1h**). Together, these developments establish a foundation for probing how head-fixed animals exhibit neural activity in response to ground motion.

### Accurate 3D reconstruction of posture

To relate neural activity with posture changes during ground motion, we recorded the animal’s 3D posture so that neural activity could be linked precisely to posture. Head-fixed mice moved their bodies in response to platform motion while we imaged under the Diesel2p mesoscope objective and captured body kinematics with four machine-vision cameras (**Fig. 2a-c**). Camera data were processed with DeepLabCut to reconstruct 3D posture (**Fig. 2d**). Accurate multi- camera geometric calibration achieved a spatial accuracy of <0.2 mm, surpassing the intrinsic accuracy of DeepLabCut. During forward tilts of the platform, the Z coordinates of the left and right hindlimb toe markers varied smoothly (**Fig. 2e**). Thus, we successfully induced a range of postures in head-fixed mice while monitoring posture with high precision.

**Figure 2.**
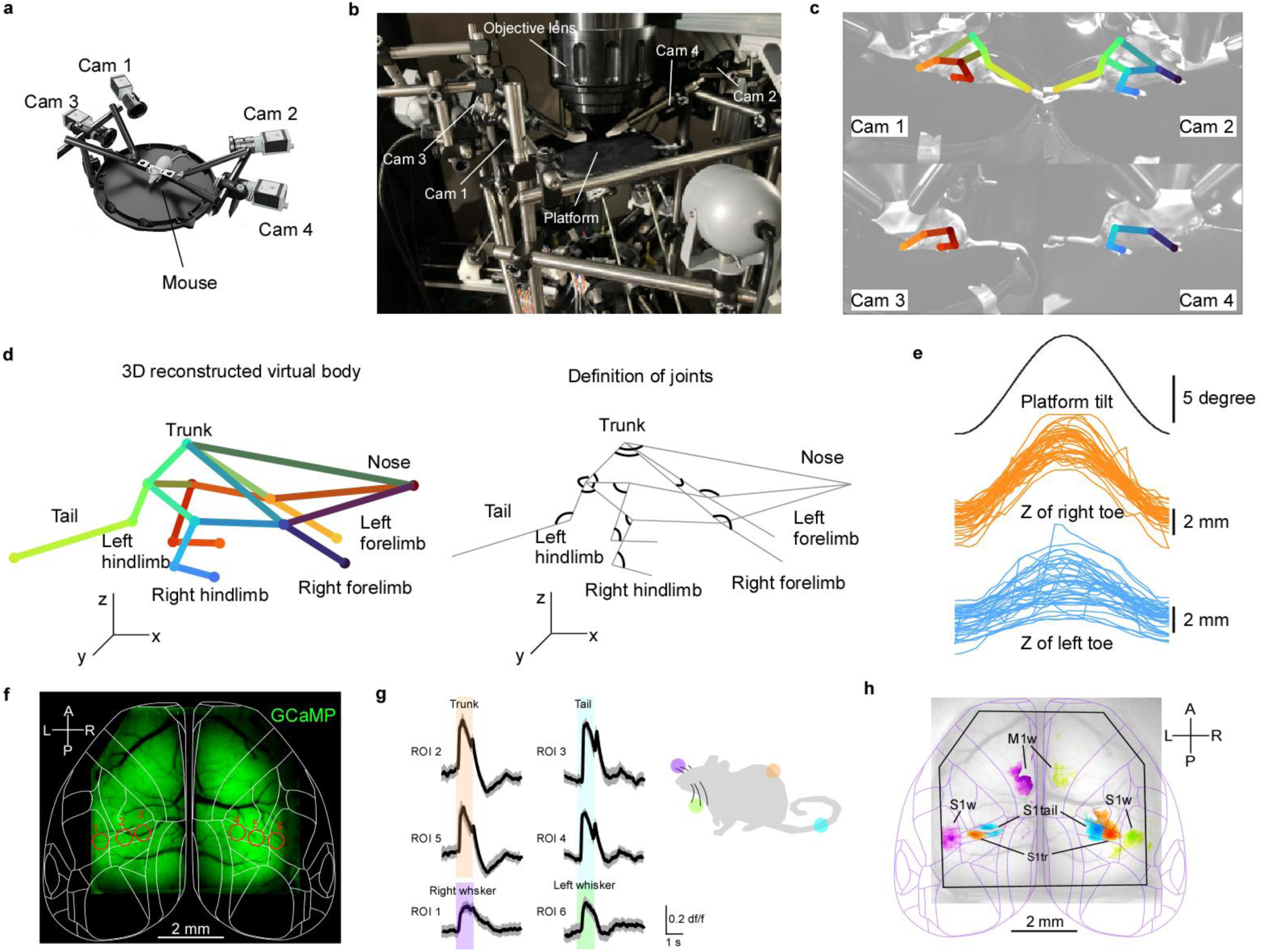
Accurate 3D reconstruction of animal posture and mapping of primary sensory cortex. **a.** Spatial arrangement of the four cameras relative to the platform and the animal. **b.** Photograph showing the positions of the platform and the four cameras placed beneath the Diesel2p objective. **c.** Mouse body imaged simultaneously by four IR-illuminated cameras during two-photon imaging; limb locations identified by DeepLabCut are overlaid. Colors correspond to panel **d**. **d.** 3D positions of 14 key points reconstructed by precise calibration of the camera coordinates (left). We also defined 16 “joints” (right) based on the key points. **e.** Changes in the Z coordinates of the left and right hindlimb toe markers while the platform tilts forward and returns. Individual traces show time courses across 40 trials. Time-aligned panels additionally show the imaging field of view (left) and the reconstructed 3D posture (right). **f.** Widefield GCaMP fluorescence through a large cranial window with overlays of dorsal cortical areas. **g.** Response of ROIs in panel f to vibration stimuli to the trunk, tail, and left or right whisker. **h.** Cortical responses to vibration stimuli to a trunk (orange), tail (cyan), right whisker (violet), and left whisker (green). S1 (trunk, tail, whisker) and M1 whisker were clearly mapped.

### Identification of sensory areas by one-photon calcium imaging

We used two preparations: mice with pan-cortical GCaMP8s expression via neonatal (“pup”) injection (n = 8), and double-transgenic mice broadly expressing GCaMP6s in cortical excitatory neurons (Ai162 × Vglut1-Cre; n = 2). To identify sensory areas, we performed one- photon calcium imaging (**Fig. 2f**). In awake, head-fixed mice implanted with a large cranial window, we delivered tactile stimulation to the trunk, tail, and whiskers and identified S1 bilaterally (**Fig. 2g**). Using these functional maps together with atlas registration, we delineated M1 (primary motor cortex), M2, S1tr (S1 trunk), S1hl (S1 hindlimb), S1fl (S1 forelimb), S1b (S1 barrel), RSC (retrosplenial cortex), and PPC in each mouse (**Fig. 2h**).

### Two-photon calcium imaging during platform motion

To broadly survey neural responses to ground tilt, we moved the platform while performing two-photon calcium imaging with a Diesel2p mesoscope. We used five motion types, including 8DT (8-direction tilt) (**Fig. 3a**). For the 8DT stimulus, the platform tilted 10° toward forward, backward, left, and right, as well as the four diagonals (front-right, front-left, back-left, back-right), and then returned to the default position. Imaging fields measured 3 × 4.25 mm^2^ and included M2, M1, S1, PPC, and RSC in either the right or left hemisphere (**Fig. 3b**). An example dataset is shown in **Fig. 3c**. We recorded a total of 20 sessions from 10 mice, yielding 6648 ± 425 active ROIs across eight cortical areas (M2: 490 ± 97; M1: 816 ± 54; S1b: 425 ± 72; S1fl: 593 ± 65; S1hl: 648 ± 77; S1tr: 464 ± 52; PPC: 902 ± 94; RSC: 1168 ± 108).

**Figure 3.**
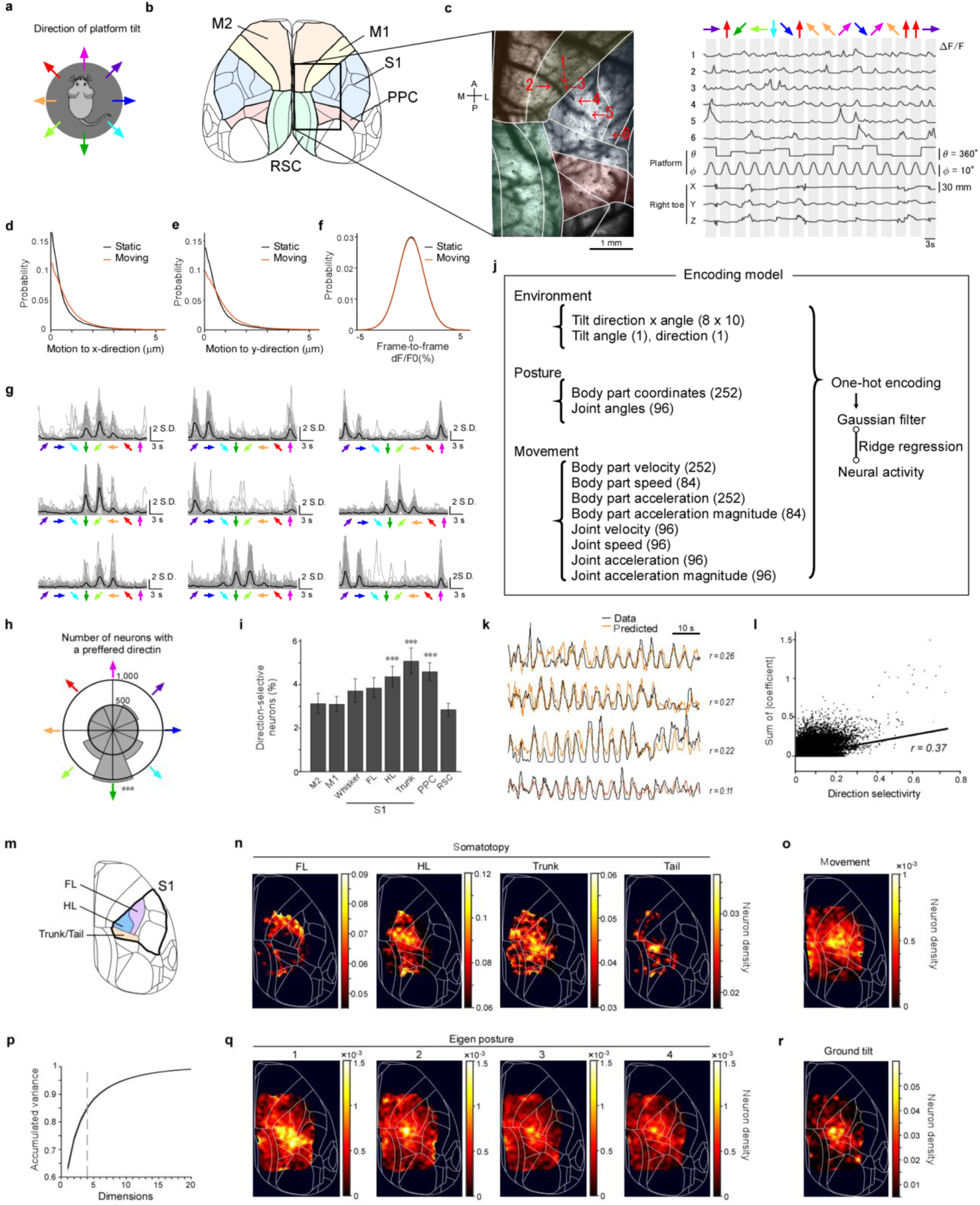
Distribution across dorsal cortex of representations related to ground tilt, movement, and posture. **a.** The platform tilted toward one of eight directions over 1 s, held for 1 s, and returned over 1 s. This 3 s sequence was repeated 40 times with directions randomized. This motion is referred to as ‘8DT’. **b.** Dorsal view of the mouse cortex and cortical areas. Rectangles indicate two-photon imaging fields, spanning M2, M1, S1, and PPC. **c.** Mean two-photon GCaMP image (left) with cortical areas overlaid in distinct colors. Right, single-neuron activity traces together with tilt direction (θ), tilt magnitude (φ), and representative body-part kinematics. Neuron IDs correspond to the cells marked at left. **d.** X-axis image drift when the platform was static versus moving. **e.** Y-axis image drift when the platform was static versus moving. **f.** Frame-to-frame fluorescence changes within each ROI during static versus moving periods, computed between adjacent time bins. **g.** Direction selectivity of 9 example neurons. **h.** Distribution of preferred directions among 4863 direction-selective neurons (χ² test, p<10^-5^; *** p<10^-5^, one-sided binomial test, Bonferroni-corrected). **i.** Percentage of direction-selective neurons by cortical area (χ² test, p<10^-3^; ***p<10^-3^, one-sided Fisher’s exact vs pooled others, Bonferroni-corrected). **j.** Schematic of ridge regression. Explanatory variables comprised posture, movement, and environmental parameters (see text). **k.** Example neuron: observed activity (blue) and ridge-regression prediction (red). **l.** Comparison between direction selectivity and the sum of absolute coefficients for direction parameters in the ridge model. A positive correlation was observed (Pearson’s correlation r=0.37, p<10^-3^). **m.** Somatotopic organization within the sensory cortex for forelimb (FL), hindlimb (HL), and trunk/tail. **n.** Map of the fraction of neurons whose activity is explained by body-part–specific parameters. The distribution aligns with locations showing one-photon responses to stimulation of the corresponding body parts. **o.** Distribution of neurons encoding movement independent of which body part was involved. **p.** PCA of posture time series (14 body parts, total 42 parameters). The plot shows the cumulative variance explained by eigenpostures from the 1st to the 20-th component (n=20 sessions). **q.** Map of neurons whose activity is best explained by the top four principal components of posture. **r.** Map of neurons whose activity is explained by ground-tilt variables. These cells were concentrated near the S1hl–S1tr border and were also distributed in PPC. Note that the peak locations for eigen postures 2 to 4 coincide with the ground-tilt map.

### Platform motion does not degrade two-photon image quality

To estimate the impact of platform motion on imaging, we compared X- and Y- axis image drift during moving versus static periods. Drift increased only slightly along both axes during motion (**Fig. 3d,e**). If Z motion was substantial, large frame-to-frame changes in ROI fluorescence would be apparent. We quantified frame-to-frame changes in ROI fluorescence (**Fig. 3f**) and found no substantial difference between moving and static periods. Because Z motion in standard calcium imaging tends to produce abrupt downward transients due to slow decay kinetics, we quantified the incidence of such events as frames in which the change in activity fell below median minus 3 × 1.48 × MAD (median absolute deviation). The event rate was 0.30 ± 0.11% during motion and 0.29 ± 0.17% during static periods, both roughly twice the 0.15% expected under a normal distribution, but with negligible difference between conditions. Thus, platform motion had minimal impact on imaging quality.

### Identification of tilt direction-selective neurons

We asked whether individual cortical neurons show selectivity for ground-tilt direction. Examining responses during the 8DT stimulus, we found neurons with sharp tuning to a specific direction (**Fig. 3g**), with backward tilts being most prevalent (**Fig. 3h**). The fraction of direction-selective neurons was significantly higher in S1tr, S1hl, and PPC than in the other areas (**Fig. 3i**). Despite regional variation, selective neurons were broadly distributed across the dorsal cortex.

### Mapping neurons responsive to posture, movement, and ground tilt

Because animals adjust posture and generate movements during ground tilt, selectivity alone cannot isolate neurons specifically encoding ground tilt from those encoding posture or movement. We therefore applied ridge regression (**Fig. 3j**) with predictors grouped into posture, movement, and environmental variables to fit each neuron’s activity. Posture variables comprised 3D coordinates of body parts and joint angles (348 variables in total). Movement variables included time derivatives (velocities), their magnitudes (speeds), and accelerations of those posture features (528 variables in total). Environmental variables comprised tilt direction and angle, together with multiple time windows relative to tilt onset (82 variables in total). The model successfully predicted the neural activities (**Fig. 3k**). Comparing direction selectivity with the sum of absolute ridge coefficients for the tilt-direction parameters revealed a moderate correlation (Pearson’s correlation, r = 0.37; **Fig. 3l**), indicating that ridge regression successfully extracted direction selectivity information while mitigating spurious correlations from posture and movement.

Using the ridge regression results, we mapped four types of responses: body-part–specific (somatotopy), movement-related, posture-related, and ground-tilt-related. First, somatotopy emerged despite the absence of explicit body-part stimulation in these experiments: parameters for forelimb, hindlimb, trunk, and tail were centered on the corresponding S1 subregions (**Fig. 3m,n**), consistent with the atlas and one-photon sensory maps obtained with vibratory stimulation. Second, neurons encoding movements were widespread, centered on S1 and extending across M1, M2, and anterior RSC, whereas PPC contained few such neurons (**Fig. 3o**).

To assess the neural encoding of whole-body posture patterns, we performed PCA (principal component analysis) on posture-related variables. Each principal component and its associated score was referred to as an “**eigenposture**,” which captures coordinated, whole-body movements (Marshall et al. 2021; Dunn et al. 2021). The top four principal components explained >80% of the variance (**Fig. 3p**), indicating that postural patterns are largely confined to a low-dimensional space. We used the scores of the top four components as explanatory variables in the ridge regression model and estimated their individual contributions. The first component was broadly distributed across S1, especially the trunk area, and extended into M1, M2, PPC, and RSC. Components 2–4 were also widely distributed, with centers near the S1hl–S1tr border. These results suggest that whole-body posture representations are shared across many areas, with hubs in trunk and hindlimb S1 (**Fig. 3q**).

Finally, neurons encoding ground-tilt variables were concentrated in S1hl and S1tr and were also present in PPC (**Fig. 3r**). Notably, the peaks of eigen postures 2–4 coincided with the ground-tilt map. Together, these findings indicate that S1hl and S1tr, while containing many neurons selective for hindlimb and trunk, also integrate ground-tilt signals with multi-joint, whole-body posture information.

### Universality of the tilt space

Does ground tilt constitute a continuous attractor that remains independent of postural and movement details, analogous to head-direction cell ensembles? To address this, we compared single-neuron responses between the 8DT and T-Rot stimuli. In three representative neurons, the preferred directions identified during 8DT were consistent with those measured under T-Rot (**Fig. 4a**). The activity of direction-selective neurons aligned to the tilt phase showed strong correlations between 8DT and T-Rot (**Fig. 4b**). These findings indicate that ground tilt is encoded in a universal format, invariant to the type of platform motion.

**Figure 4.**
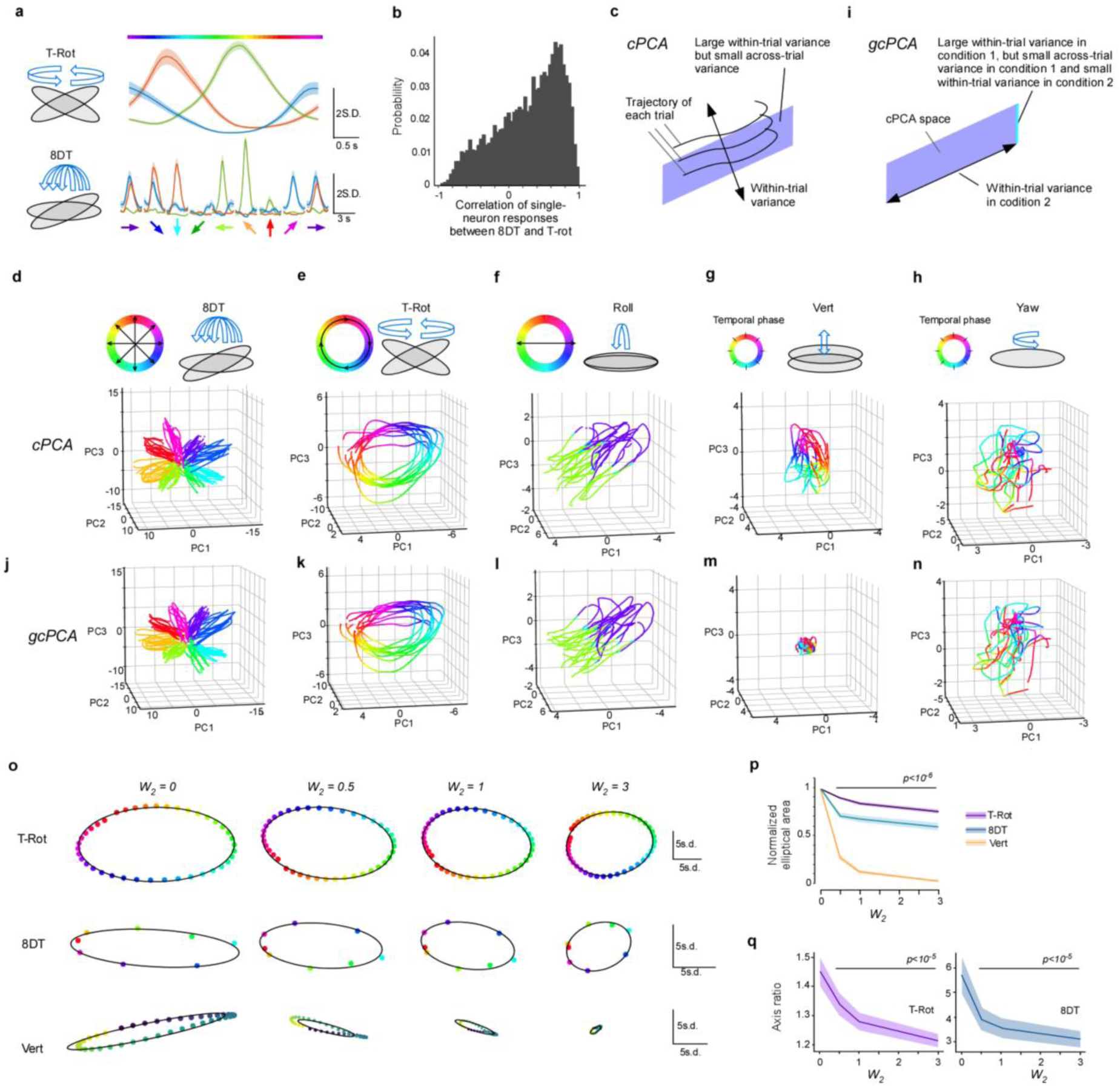
Identification of a universal representational space for tilt direction. **a.** Example activity from three neurons during tilt-rotation (top) and the responses of the same neurons during the 8DT stimulus. Direction-selective activity is preserved across the two paradigms. **b.** Correlation coefficients between phase-selective responses to T-Rot and 8DT in individual direction-selective neurons. **c.** Contrastive PCA (cPCA) defines axes that maximize within-trial variance while minimizing across-trial variance for population activity. The trade-off between these two terms is controlled by a parameter 𝑤. **d.** cPCA applied to responses during the8DT stimulus. Population activity that mirrors tilt direction appears in the top three cPCA dimensions. Color denotes tilt direction. **e.** The same neuronal population, projected onto the same cPCA axes as in **d**, but plotted for activity during T-Rot. Color denotes instantaneous tilt direction. Note that the rotation topology is preserved using axes derived from the8DT condition rather than from the T-Rot data. **f-h.** The same population and cPCA axes, plotted for a Roll session (**f**), a Vert session (**g**), and a Yaw session (**h**). **i.** Generalized cPCA (gcPCA) extends cPCA to identify axes with large within-trial variance in session 1, small across-trial variance for session 1 and session 2, and optionally reduced within-trial variance for session 2. The three-way balance is controlled by parameters 𝑤_1_ and 𝑤_2_. **j–n.** Trajectories for each stimulus type projected onto the top three gcPCA dimensions. Each of **j–n** corresponds to the panels shown in **d–h**. **o.** Representative trajectories after dimensionality reduction to two dimensions by gcPCA. 𝑤_1_ was fixed at 0.5, and four values of 𝑤_2_ (0, 0.5, 1, and 3) were tested. The top, middle, and bottom rows correspond to T-Rot, 8DT, and Vert stimuli, respectively. When 𝑤_2_ = 0 (equivalent to cPCA), ellipses were observed not only for T-Rot and 8DT but also for Vert. As 𝑤_2_ increased, the ellipse for Vert became smaller, whereas those for T-Rot and 8DT showed little change. **p.** Normalized ellipse area as a function of 𝑤_2_ , with 𝑤_1_ fixed at 0.5 (n = 20 sessions from 10 mice). For Vert, the ellipse area decreased significantly compared with 𝑤_2_ = 0, whereas the decreases for T-Rot and 8DT were significantly smaller (p<10^-6^, Per-mouse proportional slopes compared between conditions using Wilcoxon signed- rank tests, Bonferroni corrected). **q.** Same analysis as in **p**, but for the ellipse axis ratio (p<10^-5^, Wilcoxon’s signed-rank test against 𝑤_2_ = 0, Bonferroni corrected).

We next asked whether a neural manifold specific to ground tilt exists. Using cPCA (**Fig. 4c**), we maximized the following objective:

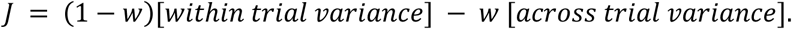

This method extracts axes that exhibit small trial-to-trial fluctuations but large task-related modulation. The cPCA analysis applied to the 8DT data revealed that activity during the 8DT traced trajectories preserving the stimulus topology, with smooth transitions among the eight directions (**Fig. 4d**). When the same axes were used to visualize responses during T-Rot, trajectories formed a circular orbit parameterized by the instantaneous tilt angle **(Fig. 4e**). Importantly, these axes were obtained from the 8DT data, not from the T-Rot data. Similar projections during Roll sessions aligned with its direction (**Fig. 4f**). These results suggest that a “tilt-space” is embedded in dorsal cortical population activity and is expressed universally across ground motion patterns.

We then projected activity during Vert and Yaw onto the cPCA space derived from the 8DT (**Fig. 4g, h**). Intriguingly, Vert also produced a ring-like trajectory, implying that this space is not a purely tilt-specific. In contrast, Yaw did not yield a ring structure. Thus, the cPCA space derived from the 8DT captures tilt-space but also partially incorporates the other components.

### Introducing gcPCA and distilling the tilt space

To suppress the Vert component and isolate the pure tilt-space, we extended cPCA and revised the objective (**Fig. 4i; Methods**):

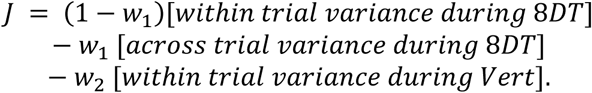

We refer to this as generalized cPCA (**gcPCA**). We visualized trajectories for all five stimulus types in the top three gcPCA dimensions (**Fig. 4j-n**). The ring-structure associated with Vert disappeared, whereas the structure for the 8DT, T-Rot, and Roll were preserved. Thus, the gcPCA demonstrated the existence of a pure tilt-space.

To examine whether the universality of the tilt-space holds statistically across different animals and sessions, we evaluated ellipses in a lower-dimensional space. We applied gcPCA using T-Rot as index 1 and Vert as index 2, and projected the data onto the principal two-dimensional plane. At 𝑤_2_ = 0, ellipses were observed for T-Rot, Vert, and 8DT (**Fig. 4o**). As 𝑤_2_ increased, the ellipse for Vert progressively shrank. When quantified by ellipse area, the reductions in area for T-Rot and 8DT were significantly smaller than that for Vert (**Fig. 4p**). Notably, the axis ratios of these ellipses decreased significantly with increasing 𝑤_2_, indicating that removal of Vert-related activity extracted the uniformity of the tilt-space (**Fig. 4q**). Thus, gcPCA revealed that the dorsal cortex represents the tilt-space in a universal manner.

### Coordinated representation of ground-tilt across dorsal cortex

Next, we compared how much information could be read out from each dorsal cortical area under each stimulus condition (**Fig. 5a**). Across conditions, readout efficiency was highest from S1tr, S1hl, and PPC. Even in areas with lower average efficiency such as M2, M1, and RSC, the scores were at least approximately 60% of those in S1tr, S1hl, and PPC. Thus, localization of information is modest, indicating that the dorsal cortex broadly and redundantly represents ground-tilt information.

**Figure 5.**
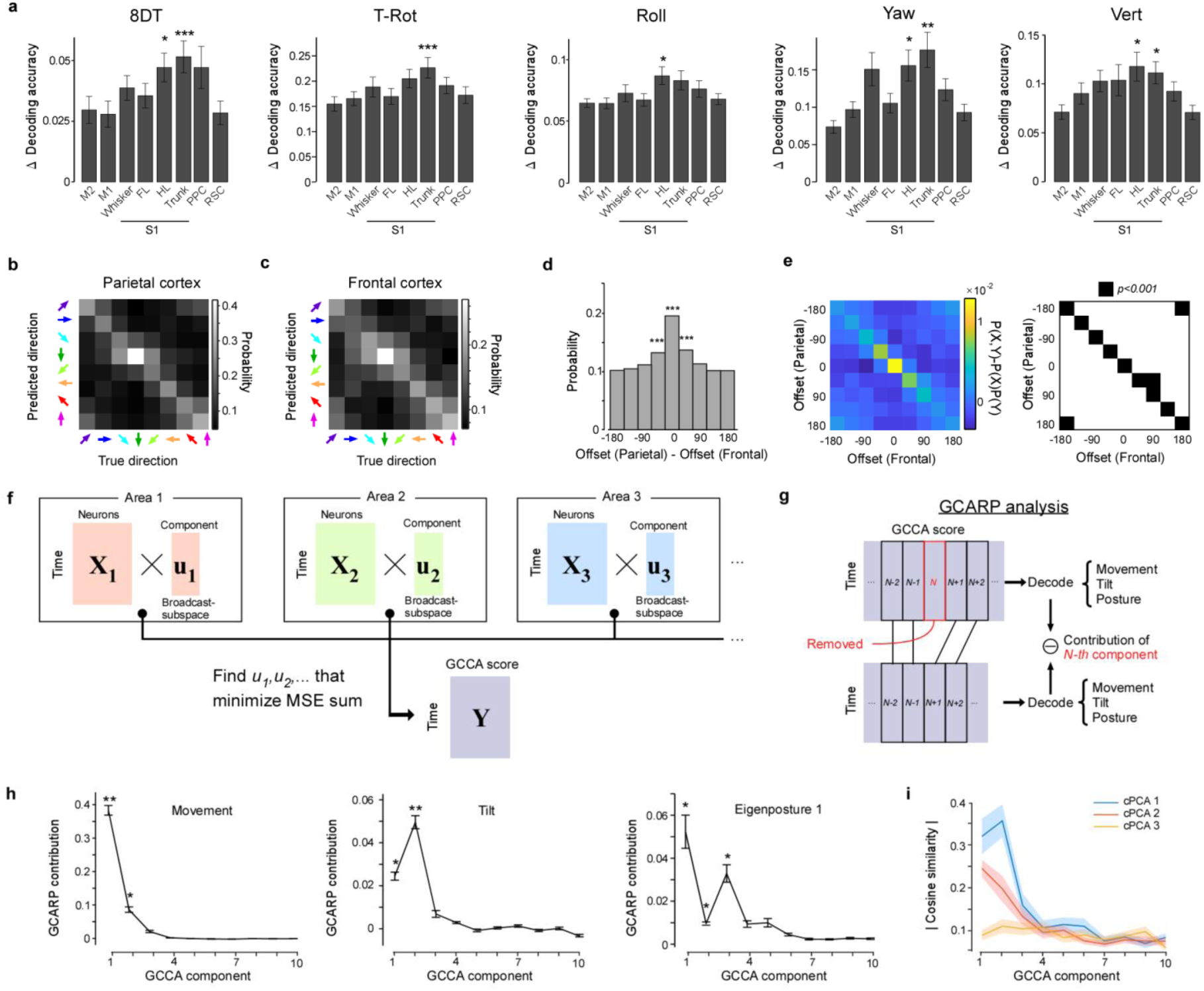
Distributed and coordinated representations of ground tilt in dorsal cortex revealed by decoding and GCARP analysis. **a.** Decoding analysis. From each cortical area, 200 neurons were randomly sampled for decoding. For T-Rot and 8DT sessions, performance was defined as the probability that the decoded tilt direction was within 45° of the true direction. For Roll, Yaw, and Vert sessions, performance was the R^2^ of the decoded parameter. In all cases, the Δscore was computed as the difference between the observed score and the mean score from randomly shuffled data. Across all motion types, Δscores for S1tr and S1hl exceeded those of other areas (*p<0.05; ** p<0.01; *** p<10^-3^; one vs rest, Wilcoxon’s rank-sum test, Bonferroni corrected). **b.** Confusion matrix of tilt-direction decoding during the 8DT session using neurons in the parietal cortex (S1 and PPC). **c.** Same analysis as in **b**, using neurons in the frontal cortex (M1 and M2). **d.** Histogram of the difference between decoded angles from frontal versus parietal cortex (from **c**). The zero bin was significantly overrepresented (*** p<10^-3^; Binomial test, Bonferroni corrected). **e.** Left, the joint distribution of decoding results from the frontal and parietal cortices after subtracting the distribution expected under independence (product of the marginals). Right, bins with significantly elevated probability are shown in black (p<10^-3^; Binomial test with Bonferroni correction). **f.** Schematic of the GCCA (generalized canonical correlation analysis). **g.** Schematic of the GCARP (generalized canonical component ablation-based representational probing). **h.** GCARP results showing shared dynamics aligned with movement (left), ground tilt (middle), and posture (eigenposture-1, right) (*p<0.05, **p <10^-3^; Wilcoxon’s signed rank test, Bonferroni corrected). **i.** Comparison of the absolute cosine similarity between the 1st–3rd components of cPCA in 8DT and the 1st–10th components of GCCA.

Given that environmental, bodily, and movement information is widely distributed across the dorsal cortex, we next asked whether these representations are coordinated. Under the 8DT condition, we performed decoding using neurons from the parietal cortex (S1 and PPC) and from the frontal cortex (M1 and M2) (**Fig. 5b,c**). Decoding using the parietal cortex alone achieved higher mutual information than the frontal cortex (parietal: 0.23 bits; frontal: 0.087 bits). To examine the coordination between the frontal and parietal cortices, we plotted the histogram of the offset difference between the two regions. The offset was defined as the difference between the true direction and the decoded direction. The offsets were significantly aligned, indicating coordinated representation of tilt direction (**Fig. 5d**). Furthermore, the joint distribution of decoding offsets in the frontal and parietal cortices showed that the diagonal elements, representing the agreement between the two cortices, were significantly higher than the probability expected under independence (product of the marginals) (**Fig. 5e**). This indicates that even when both the frontal and parietal cortices produced errors, they nevertheless exhibited coordinated representations. Together, these results demonstrate that the dorsal cortex constructs an internally consistent ground-tilt representation through functional coordination.

### Broadcast-subspace analysis (“GCARP”) reveals the information content of shared dynamics

Signals related to movement and arousal are known to dominate wide cortical activity during spontaneous behavior and decision-making tasks (Stringer et al., 2019; Musall et al., 2019). To examine the information content of the shared cortical dynamics, we recently introduced CARP (Canonical Component Ablation-based Representational Probing) (Imamura et al. 2025), which can quantify the representation content within the communication subspace (Semedo et al., 2019). Here, we extend this approach from two areas to three or more by replacing CCA with generalized CCA (**GCCA**). We refer to this extended method as **GCARP** (generalized CARP), and to the subspace extracted with GCARP as the **broadcast-subspace**. The broadcast-subspace is maintained by each individual cortical area and contains information that is shared not between pairs of areas but across all areas examined.

GCCA applies a linear transformation to the population activity of each area and extracts latent time series that are maximally correlated across all transformed series (**Fig. 5f**). The resulting axes are termed canonical components, and their trajectories are referred to as canonical scores. These components are ordered according to the extent of shared variance across areas. In GCARP, we first decoded target variables from the full set of canonical scores, which represent the common dynamics expressed across all recorded areas. Decoding based on these scores therefore captures information jointly represented across areas. We then ablated one component by excluding its score and repeated decoding with the remaining scores. The reduction in decoding performance quantifies the contribution of the removed component to the target variable, thereby revealing which shared components across areas carry which types of information (**Fig. 5g**).

In the 8DT session, we applied GCARP to neural activity recorded from eight areas (M2, M1, S1b, S1hl, S1fl, S1tr, PPC, and RSC) and evaluated three types of information: movement, ground tilt, and eigenposture (the first component). The significant axes were the 1st–2nd GCCA components for movement and tilt, and the 1st–3rd GCCA components for posture. Interestingly, movement was most strongly represented in the first GCCA component, whereas ground tilt was most strongly represented in the second, and posture was also significantly represented in the third (**Fig. 5h**). Thus, the dynamics shared across wide regions of the dorsal cortex reflect movement, tilt, and posture, and exhibit a clear structural organization.

### Relationship between tilt-space and shared cortical dynamics

GCARP analysis demonstrated that the shared dynamics contained tilt information. To more directly examine the relationship between the shared dynamics and the tilt-space, we compared the top three dimensions of cPCA with the 1st–10th GCCA components using cosine similarity. The strongest alignment was observed between the first cPCA component, which most strongly represented tilt direction, and the second GCCA component, which GCARP had identified as carrying tilt information (**Fig. 5i**). Thus, the broadcast-subspace robustly captures tilt information, as confirmed both by GCARP and by direct comparison of the subspaces.

## Discussion

By combining a newly developed 6-DoF platform, 6AGM, for head-fixed mice with a Diesel2p mesoscope, we simultaneously imaged neural activity from dorsal cortical areas including M1, M2, S1tr, S1hl, S1fl, S1b, PPC, and RSC, together with 3D postural kinematics at high spatiotemporal resolution during ground-tilt stimulation. Analyses of neurons related to movement, posture, and the environment (tilt) revealed that (i) neurons in the dorsal cortex exhibit sharp selectivity for tilt direction (“direction-selective neurons”); (ii) their activity forms a “tilt-space” that preserves a circular phase independent of the method of tilting; and (iii) broadcast-subspace analysis, GCARP, revealed that movement, tilt, and posture information were shared in the top three GCCA dimensions. These findings suggest a basic principle whereby the cerebral cortex extracts environmental information in specialized cortical circuits, while distributing and integrating it across a variety of cortical areas to enable control of posture and movement.

The localization of tilt-selective neurons at the S1tr–S1hl border indicates that these regions function as “hubs” that integrate multi-joint inputs and extract environmental geometry. The same neurons maintained consistent preferred angles across eight tilt-direction conditions and under rotational stimulation, and rotation trajectories traced a continuous ring in cPCA space. We therefore propose that tilt information is represented in a “universal coordinate system” that is independent of postural details, ongoing movement, and mechanical load. Rather than reflecting extraction of elemental features in the environment, this appears to be an abstract representation assembled by mainly integrating limb-trunk information. In rats, the border region between the rat PPC and S1 shows a less distinct layer 4 compared to other parts of S1, exhibiting a cytoarchitecture closer to that of association areas (Olsen and Witter 2016; Haghir et al. 2024). Therefore, at least in rodents, the S1tr-S1hl differs from other primary sensory cortices in that it can engage in more integrative information processing.

The continuous phase representation of tilt may constitute an attractor for a “tilt-space” variable, comparable to the ring-attractor of head-direction cells (Peyrache et al. 2015), the 2D continuous attractors of place cells (McNaughton et al. 2006; Yoon et al. 2013), and the toroidal attractor of grid cells (Gardner et al. 2021). Although head-direction cells are commonly observed in RSC (Chen et al. 1994), tilt-related signals were not specifically localized to RSC but S1 in our experiment. This suggests that along the gradient from hippocampal structures supporting abstract spatial representations to early sensory cortices processing egocentric bodily information, ground-tilt information resides relatively closer to the sensory end, which is unsurprising given that it is a parameter directly involved in motor execution. It is also consistent that PPC represents ground-tilt information, as it can provide spatial context to visual and auditory streams (Yao et al. 2020; Licata et al. 2017). Stabilizing posture requires the relative angle between gravity and the ground normal; by coupling tilt-space with head direction and visual/vestibular inputs, the brain could rapidly estimate the energetic costs necessary for whole-body coordination and route selection.

Tilt-selective neurons were most abundant at the S1tr–S1hl border, as predicted by previous studies (Nandakumar et al. 2021; Disse et al. 2023), but they were also scattered across other dorsal cortical areas, indicating that the tilt-space itself is distributed throughout the cortex. To investigate the dynamics broadly shared across the cortex, we developed GCARP. According to GCARP, the first component of the broadcast-subspace primarily captured movement variables, whereas the tilt-space aligned with the second component. The identification of the first component as a “movement axis” is consistent with our previous findings (H. Imamura et al., 2025). Shared posture information was also prominently represented in the third component. Taken together, these results indicate that the broadcast- subspace integrates both dynamic and static body information as well as information about the peri-personal environment. Thus, the essence of dynamics within the broadcast-subspace may lie in forming an integrated body–environment representation that underlies diverse forms of information processing in the animal.

Our fully open-source six-degree-of-freedom platform (6AGM) provides the stability required for two-photon imaging under head fixation while allowing arbitrary translations and rotations. Mounting a treadmill on the platform would enable navigation experiments involving slopes. Compared with floating balls (Dombeck et al. 2007), rotating disks (Hoogland et al. 2015), or wheels (Najafi et al. 2014), this setup provides a more natural environment and substantially expands the degrees of freedom for navigation paradigms. Coupling the system to a head-rotation module (Voigts and Harnett 2020) may further allow free exploration of two-dimensional spaces with slopes. By employing this platform to investigate brain disease models with postural instability, progress may be achieved in pathophysiological research, therapeutic development, and rehabilitation, ultimately contributing to improved human quality of life. Thus, this platform constitutes foundational infrastructure with broad applications, ranging from basic neuroscience to clinical neurology.

## Materials and methods

### Animal preparation

All experiments were approved by the Animal Research Ethics Committee of the Institutional Animal Care and Use Committee of Tokyo Medical and Dental University (A2021-290C6, A2024-060C2) and were carried out in accordance with the Fundamental Guidelines for Proper Conduct of Animal Experiment and Related Activities in Academic Research Institutions (Ministry of Education, Culture, Sports, Science and Technology of Japan). Two double-transgenic adult mice expressing GCaMP6s and eight adult C57BL/6Jcl mice expressing GCaMP8s through neonatal AAV (adeno-associated virus) injection were used. Double-transgenic mice (C57BL/6JJcl, male or female) were made by crossing the Vglut1-Cre mice and Ai162 mice (Daigle et al. 2018) that can express GCaMP6s in excitatory neurons in a Cre-dependent manner. Doxycycline was administered to Vglut1-Cre and Ai162 mice during development to suppress transgene expression and mitigate potential developmental effects of calcium buffering.

### Neonatal AAV injection

Eight mice in the study underwent neonatal virus injection on postnatal day 1 or 2 (Oomoto et al. 2021). The pups were placed on an ice bed for anesthesia. A custom head-fixation device developed by the NARISHIGE group was used to fix the pup’s head. A borosilicate glass pipette with an outer diameter of 50 μm and inner diameter of 30 μm was inserted into the head 250–300 μm deep from the surface. Four μl of AAVDJ-Syn-jGCaMP8s (Y. Zhang et al. 2023) at a titer of 1.16 × 10^13^ was injected over a period of 1 minute. The process was repeated for both hemispheres for 2 mice. The pups were then placed on a recovery box on top of a healing mat to promote recovery. The entire procedure took no more than 10 minutes per pup.

### Head plate surgery

Mice were anesthetized with isoflurane (1-1.5%) and the body temperature was monitored with a rectal thermometer. Eye ointment was applied to keep the eyes moist. Head plate surgery was conducted on mice over 8 weeks of age. A custom head-fixing imaging chamber (Hira et al., 2013) was mounted on the skull with a self-cure dental adhesive resin cement (Super bond, L-Type Radiopaque; Sun Medical) and luting cement (FUJI LUTE BC; GC Dental). The ample width of the head plate allowed the implantation of a 6 mm × 6 mm glass plug and subsequent large field-of-view two-photon imaging. The surface of the intact skull was then coated with clear acrylic dental resin (Super bond, Polymer clear; Sun Medical) and silicon to prevent drying.

### Cranial window implant

Three days after head plate surgery, an ultra-large cranial window implant (Hattori & Komiyama, 2022) was performed as follows. The glass plug used to cover dorsal cortex was made with a 6 mm × 6 mm #3 cover slip and a 7 mm × 7 mm #0 coverslip which were custom made by Matsunami Glass Ind. Two coverslips were glued together using optical adhesive (Norland Optical Adhesive 63; Norland Products). After anesthesia, 4 μl/g body weight of 20 % mannitol solution was administered via intraperitoneal injection to reduce brain swelling, and 0.216 μg/body weight (g) of atropine sulfate was administered intraperitoneally to reduce saliva and mucus secretion. The mouse head plate was fixed after anesthesia. Clear acrylic dental resin was removed using a dental drill, followed by marking of bregma and stereotaxic coordinates. The skull along the outline of the cranial window as well as coronal and sagittal sutures was thinned with the drill. The frontal and parietal bones were removed in four separate pieces. The glass plug was then placed on the exposed cortex, enabling optic access to the entire dorsal cortex. The glass plug was attached to the bone with cyanoacrylate glue and secured using dental resin cement.

### Two-photon imaging

Two-photon imaging was conducted with a custom-made mesoscope, Diesel2p (Yu et al. 2021; Hira et al. 2025). For excitation of GCaMP, we used a fixed-wavelength fiber laser, ALCOR 920-2 (920 nm, 80MHz). The imaging was done on two FOVs, each with size 4.25 ×1.5 mm^2^, totaling an FOV of 4.25 × 3 mm^2^. The number of pixels of the image was 1,500 by 1,168 and images were sampled at 4.79 frames per second.

### One-photon calcium imaging

One-photon calcium imaging was conducted more than two days before or more than three days after two-photon calcium imaging on the platform. Mice were imaged with head-fixed conditions. Excitation light for GCaMP was delivered in a dark enclosure. A 470 nm blue LED was positioned 15° above the right hemisphere. Emitted light passed an 85 mm F/2 lens, a 50 mm F/1.4 lens, and a 525/39 nm filter before reaching a CMOS camera (504 × 504 pixels, 14.1 µm pixel size after 3 × 3 binning). Frames were acquired at 20 fps under LabVIEW control. The recording was conducted in an awake state and lasted less than 30 min.

### Sensory stimulation

For sensory stimulation of mice, a plastic pipette was attached to the membrane of a speaker. The pipette was then attached to either the tail, trunk or right/left whisker of the head-fixed mouse. In each 8-s trial, there was a 50 Hz stimulation for 0.8s, and each body part was stimulated for 60 trials. The activity of the dorsal cortex was recorded through one-photon calcium imaging. For three mice, the location of peak response to each stimulation was mapped and aligned with the mouse brain atlas. For two mice, the dorsal cortex and the brain atlas were aligned using stereotaxic coordinates identified during cranial window implant surgery.

### Image processing

Image preprocessing was done using ImageJ (National Institutes of Health) and MATLAB (MathWorks,Natick, MA, USA) software. The bidirectional phase offset of the image was calculated, and sinusoidal distortions caused by the resonant scanner were corrected. Turgoreg plugin (Thévenaz, Ruttimann, and Unser 1998) on ImageJ was used for image registration. By utilizing ImageJ macros, the images were divided into 9 by 8 smaller blocks, and registration was performed for each block. Subsequently, MATLAB was used to combine the drift-corrected images from each block. The X- and Y- offsets of each block was linearly interpolated across all of the pixels to ensure consistency of the shift. The registered images were then analyzed using Suite2p (Pachitariu et al. 2016) to determine the location of the regions of interest (ROIs). The intensity of each ROI was calculated by subtracting mean values of background pixels within 120 μm of the ROI excluding the other ROIs from mean value of the ROI. For each ROI, the 8th percentile pixel value within 15 s of the time window was obtained, which was then smoothed using a Gaussian filter with standard deviation of 30 s. This was considered as the baseline of neural activity, and was subtracted from the original data to set the baseline to 0. A Wiener filter with temporal length of 1.7 s was applied to reduce noise. Non-negative deconvolution with a time constant of 0.48 s for GCaMP8s and 1.2 s for mouse GCaMP6s (fast-oopsi; Vogelstein et al. 2010) was performed, and a Gaussian filter with standard deviation of 0.2 s was applied at the end. We defined the time-series data for each ROI as its “activity.” ROIs whose activity skewness exceeded 0.5 were classified as active neurons (or simply “neurons”), and only these were included in analyses. For analysis, the neural data was interpolated to have a 10 Hz signal rate.

### Platform motion during two-photon imaging

We delivered five types of platform motion to each animal during two-photon calcium imaging.

1. **8DT (8-direction tilt):** Each trial began with a 1-s baseline, followed by a 10° tilt over

1.5 s toward one of eight possible directions, and then a 1.5-s return to baseline. To prevent the mouse from predicting the upcoming tilt direction, the sequence was constructed so that every possible pair of consecutive directions occurred exactly once while maintaining equal presentation of all directions. This required 65 trials in total from which the first trial was omitted for analysis, resulting in 64 trials (8 per direction). The tilt trajectory was generated using a cosine profile to provide smooth acceleration and deceleration.

1. **T-Rot (Tilted-rotation):** The platform was tilted 10° forward and, while maintaining this tilt, rotated clockwise for 10 full revolutions (4 s per revolution). Each session therefore comprised 10 trials, but the first trial was omitted from analysis, leaving 9 trials in total.
2. **Roll (left–right tilt):** The platform was initially tilted 10° to the left and then seesawed back and forth between left and right. Each cycle lasted 4 s and was repeated 10 times. To ensure smooth acceleration and deceleration, the tilt angle followed a cosine profile. As with the other motion types, the first trial was omitted from analysis, leaving 9 trials in total.
3. **Yaw (horizontal rotation):** The platform rotated about its central vertical axis by ±15° and oscillated back and forth. Each cycle lasted 4 s and was repeated 10 times. The rotation followed a cosine profile to ensure smooth acceleration and deceleration. As with the other motion types, the first trial was omitted from analysis, leaving 9 trials in total.
4. **Vert (vertical translation):** The platform translated vertically between +5 mm and −5 mm in an oscillatory motion. Each cycle lasted 4 s and was repeated 10 times. The translation followed a cosine profile to ensure smooth acceleration and deceleration. As with the other motion types, the first trial was omitted from analysis, leaving 9 trials in total.

Each session was composed of repeated *motion blocks*, with each block consisting of T-Rot (10 trials), Roll (10 trials), Yaw (10 trials), Vert (10 trials), and 8DT (65 trials). Each block was repeated 6–8 times within a session, resulting in a total session duration of 42–56 minutes. Neural activity was stably recorded during these platform motions, allowing us to sample the same neuronal population across conditions.

Details of the newly developed platform (capable of generating arbitrary ground motions including those above), its operating principles, components and hardware, software, motion parameters, and comparisons with existing systems are provided in the **Supplementary Information**.

### Habituation

In order to habituate animals to ground motion under head-fixed conditions, we placed head-fixed mice on a balance board with a diameter of 250 mm (HCF-BDBUL, Elecom) and randomly tilted the board for 30 minutes. The board was perturbed by a rotating zip tie attached to a stepper motor, causing random tilts, and the head-fixed mice had to maintain their balance at the center of the board. Four mice went through this habituation stage for 4 days, followed by a phase of habituation to movement on a 6DOF platform under a microscope, after which the imaging experiment was performed. Six mice were habituated directly on the 6AGM after head plate fixation habituation.

### Reconstruction of 3D posture

Following the removal of hair from the mouse’s limbs and back, infrared absorptive marker was applied to their elbows, knees and three equally spaced points on the back. Four cameras were used to record mouse movements, and DeepLabCut (Nath et al. 2018) was used to track left and right paws, left and right elbows, left and right toes, left and right heels, left and right knees, a point of the tail, and the three points on the back (14 points in total; **Fig. 2d**). The nose position was assumed to be a single fixed point because it does not move during head fixation. The detected points were reconstructed to 3D coordinates by triangulation. Any frames with triangulation error >4 mm were replaced by interpolated values derived from neighboring frames with error <4 mm. Velocities and accelerations were then computed from the 3D trajectories after applying a 10 Hz low-pass filter.

For a precise triangulation, radial lens distortion correction, intrinsic parameter estimation, and stereo calibration were all performed using checkerboards (see **Supplementary Information** for detail). Following radial lens distortion estimation using two-parameter division model, 5 parameters of the pinhole camera model were linearly estimated using undistorted images. The RANSAC algorithm (Fischler and Bolles 1981) was employed to filter out inaccurately detected checkerboards, and the parameters were subsequently optimized interactively. Stereo calibration was performed using a normalized 8-point algorithm, similarly employing the RANSAC algorithm to filter out faulty checkerboards. The entire algorithm, with the exception of the checkerboard detection, was developed and customized in MATLAB.

It is not possible to measure the actual joint angle of an animal from our body part estimation. Therefore, we adopted an angle consisting of the estimated body parts as the “Joint” (**FIg. 2d**). Since an angle can be determined once three points are determined, 14C3 = 364 angles can be selected from the 14 estimated body part points. We used 16 of these angles as joints, and the results did not change when all 364 angles were used to estimate neural activity.

### Direction selectivity

Tilt direction selectivity of individual neurons was obtained from the data during 8DT motions as follows. For each neuron, the mean activity during the baseline period, defined as the horizontal condition without ground motion, was calculated. For each direction, the mean activity during 1–2 s after stimulus onset was measured, and the baseline activity was subtracted. Negative values were set to zero. The response for each direction was then normalized by dividing by the sum of responses across all directions. The direction selectivity score was defined as the magnitude of the vector sum of the normalized responses, where the normalized response for each direction was treated as the vector magnitude. The preferred direction of each neuron was defined as the direction that yielded the maximum response. Neurons with high direction selectivity were defined as those with a direction selectivity score >0.1 and a significant effect of direction, assessed by one-way ANOVA across tilt directions for each neuron (p <0.05).

### Encoding model

The encoding model was implemented as a ridge linear regression that explained the activity of individual neurons using environmental, postural, and movement parameters. These parameter sets (total 1,486 parameters) were defined as follows.

- Environmental information comprised tilt direction and tilt angle, and they were jointly discretized using 8 and 5 bins, respectively, yielding 80 bins in total. The bins were one-hot encoded and were temporally filtered with a Gaussian kernel (σ = 0.1 s). In addition to the one-hot vectors, the non-discretized tilt direction and tilt angle were included, yielding a total of 82 environmental parameters.
- Postural information consisted of the 3D coordinates of 14 body parts and the joint angles computed from these positions. The 3D coordinates were discretized into five bins individually and represented as one-hot vectors, yielding 14 (body parts) × 3 (XYZ) × 5 (bins) = 210 parameters. Each joint angle was discretized into five bins and one-hot encoded, giving 16 × 5 = 80 parameters. All one-hot–encoded time series were temporally filtered with a Gaussian kernel (σ = 0.1 s). Non-discretized parameters were also included, resulting in a total of 348 postural parameters, 252 parameters for the 3D coordinates and 96 for the joint angles.
- Movement information included, for each of the 14 body parts, velocity and acceleration along the XYZ axes, as well as the magnitudes (speed and acceleration magnitude). Each parameter was discretized into five bins and represented as one-hot vectors. Velocities across body parts comprised 252 parameters, whereas velocity magnitudes, which are scalars over XYZ, comprised 84 parameters, including non-discretized values. Similarly, accelerations and its magnitude comprised 252 and 84 parameters, respectively. The joint angular velocity, acceleration, speed, and acceleration magnitude were also discretized into five bins and one-hot encoded, yielding 96 parameters each. As above, all one-hot–encoded time series were temporally filtered with a Gaussian kernel (σ = 0.1 s). A total of 1,056 movement parameters were obtained.

In the full encoding model, all parameters were used to fit the activity of individual neurons. Prediction accuracy was quantified by the coefficient of determination (R²) using 10-fold cross-validation.

- To quantify body-part–specific information, we constructed reduced models excluding parameters for each group: forelimb (paw and elbow), hindlimb (toe, heel, knee), trunk, and tail. The decrease in R² relative to the full model was taken as the contribution of each body-part group to the neuronal representation.
- To quantify movement-related information, we constructed models excluding all velocity, acceleration, and their magnitudes for all body parts and joint angles. The reduction in R² relative to the full model was defined as the representation of movement information.
- To quantify whole-body postural patterns, we performed principal component analysis (PCA) on the time series of all body-part positions and joint angles to identify eigen postures. Using all components except a given target dimension, we reconstructed the body-part and joint-angle time series and refit the model. The decrease in R² relative to the full model was defined as the representation carried by the excluded eigen posture.
- Finally, to quantify the encoding of ground tilt, we fit models that excluded tilt parameters while including all other parameters and evaluated prediction accuracy by R². The difference from the full model’s R² was taken as each neuron’s representation of ground-tilt information.

### cPCA and gcPCA

To extract the informative dimensions from the neural data, we used cPCA (Abid et al. 2018; Imamura et al. 2025) or generalized cPCA (gcPCA). cPCA finds axes that maximize the following cost function,

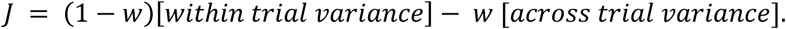

For example, in the case of 8DT, since there are eight conditions, we obtained a time series of 8 conditions × 3 seconds each, yielding 24 seconds of data, repeated over ∼50 trials, resulting in a 24 × 50 matrix. The variance computed after averaging across trials is defined as the within-trial variance, whereas the variance computed after averaging across the time axis is defined as the across-trial variance.

Parameter 𝑤 determines the balance between the two. We set 𝑤 = 0.5, unless otherwise noted. gcPCA finds axis that maximize the cost function 𝐽, defined as

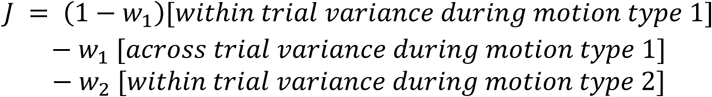

Parameter 𝑤_1_ and 𝑤_2_ determines the balance between the three variances.

### Decoding analysis

We decoded tilt-related variables (8DT, T-Rot, Roll), vertical position (Vert), and horizontal rotation direction (Yaw). All decoding analyses used bagged decision trees (MATLAB function *TreeBagger*). The number of trees was fixed at 100. For 8DT and T-Rot, tilt directions were discretized into eight categories spaced 45° apart, and classification was performed to decode the direction. For Roll, Vert, and Yaw, continuous decoding was carried out using regression. To prevent overfitting, we performed 10-fold cross-validation and recorded classification accuracy. When comparing decoding performance across regions, we randomly sampled 200 cells from the target region, reduced the data to 20 dimensions by PCA, and then performed decoding. When decoding with all areas combined, and for the frontal cortex and parietal cortex analyzed separately, we used all available cells, reduced them to 1000 dimensions by PCA and to 100 dimensions by cPCA (w = 0.5), and then decoded. In GCARP, decoding was performed on the 100-dimensional or (100−1) - dimensional data after GCCA. Because dimensionality reduction must be integrated with cross-validation, the dimensionality-reduction model was fitted on each training fold and the resulting transform was applied to the corresponding test data. We set 𝑤 = 0.5 for cPCA.

### GCCA (Generalized Canonical Correlation Analysis)

To extract latent components shared across eight cortical areas, we employed Generalized Canonical Correlation Analysis (gCCA). gCCA extends conventional canonical correlation analysis (CCA), which maximizes correlation between two datasets, to multiple datasets, allowing estimation of a common low-dimensional representation across several brain regions.

Neural activity matrices from each area 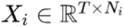, where 𝑇 is time and 𝑁_𝑖_ is the number of neurons) were used as input. Each dataset was z-scored, and dimensionality was reduced to 100 using principal component analysis (PCA). The goal of gCCA was to identify a shared latent space 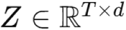 across all areas. Linear projection matrices 𝑊_𝑖_ were introduced for each area, and the optimization problem was defined as:

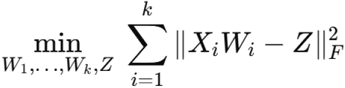

where 𝑘 = 8 is the number of areas and ‖·‖ denotes the Frobenius norm. This formulation estimates projections such that the transformed time series from all areas are aligned to the common latent representation 𝑍.

### GCARP (Generalized Canonical component-ablation based representational probing)

GCARP is an extension of CARP to multi-area datasets (Imamura et al. 2025). 200 neurons were randomly sampled from each area, which were then reduced to 100 dimensions using PCA. The dimensionally reduced time series from all areas were then transformed into a 100- dimensional representation using GCCA. Using this representation, decoding of movement, ground tilt, and eigen-posture was performed in the same manner as described above. Movement was defined as the mean speed of all the body parts. Ground tilt was decoded through classification after dividing them into 8 bins, 45° apart. The decoding accuracy obtained here was regarded as the maximal decoding performance. Next, the *N*-th GCCA dimension (N=1,2,…,10) was removed, and decoding was repeated with the remaining dimensions. The reduction in decoding accuracy relative to the maximal performance was interpreted as the contribution of the *N*-th component.

### Statistics

Student’s t-test, Wilcoxon’s rank-sum test, Wilcoxon’s signed-rank test, Spearman’s rank correlation coefficients were used for pairwise comparison. Bonferroni correction was applied for multiple comparisons. For preferred direction distribution, overall deviation from uniformity was tested with a χ² goodness-of-fit test, followed by one-sided exact binomial tests with Bonferroni correction. To compare the ratio of direction-selective neurons across cortical areas Pearson’s χ² test of independence followed by one-sided Fisher’s exact tests with a Bonferroni correction was used. Data was expressed as means ± s.e.m. unless otherwise noted.

## Acknowledgements

We thank S. Kato and K. Kobayashi for AAV production, H. Mori, S. Oota, O. Ooishi and H. Kasai for discussions, all other members in the Isomura lab for supporting this work. This work was supported by JP22wm0525007 (RH), JP19dm0207089 (YI) from AMED, JP22H02731 (RH), JP20K22678 (RH), JP21B304 (RH), JP21H05134 (RH), JP21H05135 (RH), JP21H05242 (YI), and JP23H02589 (YI) from MEXT/JSPS, JPMJCR1751 (YI) from JST, Nakatani Foundation (RH), Shimadzu Foundation (RH), Takeda Science Foundation (RH), The Precise Measurement Technology Promotion Foundation (RH), Tateishi Science and Technology Foundation (RH), and Research Foundation for OptoScience and Technology (RH).

## Author contributions

F.I. Riichiro H. conceived the project. F.I. H.I. conducted experiments and data analysis. F.I. designed and manufactured the 6AGM. Reiko H. conducted pilot experiments. Y.I. supervised the project. Riichiro H. F.I. wrote the paper with the comments of all authors.

