## Supplementary information for "Ground tilt representation in the rodent cerebral cortex"

  1.1. Platform development and comparison with existing 6-DOF platforms

  1.2. Mechanical design of 6AGM (CAD)

  1.3. Platform and stepper motor position coordinate conversion

  1.4. Discrete-time feedback–feedforward control scheme

1.5. Control system of 6AGM
1.6. Encoder interface
1.7. Actuator driver interface
1.8. Motion generation
1.9. LabVIEW software modules (Sub-VIs)

2. Camera calibration
2.1. Single camera calibration
2.2. Stereo camera calibration

### 1. 6AGM

We developed a fully open-source six-degree-of-freedom arbitrary ground-motion platform (6AGM) for use in this study and with future integration into virtual-reality (VR) systems and brain–machine interfaces (BMIs) in mind. In the sections that follow, we provide a detailed account of the platform requirements and comparisons with existing devices, the mechanical design of 6AGM, the computational methods, and the hardware and software.

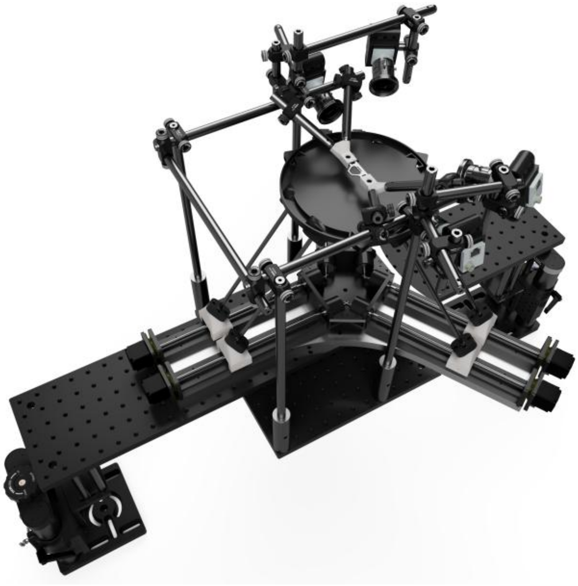

##### 1.1. Platform development and comparison with existing 6-DOF platforms

We searched for a mechanism capable of rapidly moving a platform to arbitrary positions and orientations to enable an integrated understanding of movement, posture, and environmental information using large-scale two-photon calcium imaging. For use with mice under the Diesel2p two-photon microscope, the static design constraints were an overall footprint ≤ 500 mm in diameter and ≤ 400 mm in height, with a platform disk 150–250 mm in diameter. The dynamic requirements were translational travel ≥ 60 mm in both mediolateral and anteroposterior directions, and rotational travel ≥ 10° in roll, pitch, and yaw. Because the head-fixation apparatus, lick port, speakers, and visual-stimulus monitor together weigh ~6 kg, the payload capacity needed to be ≥ 6 kg. While meeting these specifications, the system also had to be stable enough for two-photon imaging—specifically, under head fixation the brain's vertical (z-axis) motion had to be continuously suppressed to ≤ 2 μm—and we had to avoid imparting unwanted stimuli to the animal caused by jitter or chatter arising from feedback delays and related effects. We also required that arbitrary motion trajectories be programmable via custom scripts and that real-time motion be monitored with immediate response to real-time motion targets to enable future closed-loop control.

**Supplementary Table 1** summarizes examples of commercially available 6-DOF platforms. While some products met individual requirements for size, payload, travel range, or speed, none satisfied all of them simultaneously. In particular, compact units had limited maximum speeds, whereas platforms which can move fast enough were too large to mount on an optical table. Several products offered precision far beyond our needs; however, devices that prioritized speed over precision were largely simulator rigs and were very large. Therefore, we built a platform that meets these requirements in-house.

Here, we developed a fully open-source system that satisfies the above specifications. Components can be custom built, and we have released the corresponding CAD files. The hardware is based on low-cost, widely available controllers (Arduino, Arduino MEGA, and NI USB-6001), and the software centers on a LabVIEW-based control stack. All scripts are released as open source.

| Product | Maximum velocity | Range of motion | Precision(resolution / repeatability) | Size (Height, base diameter) | Mass/ load capacity | Interface/ controller |
|---|---|---|---|---|---|---|
| 6AGMv1, F. Imamura et al. | 290mm/s (X,Y), 140mm/s(Z), 90°/s(Rx), 105°(Ry), 157°(Ry) | 165/165/115 mm / ±10°/±15°/±15° | Joint resolution 7.5 μm | Height 250mm, Ø300mm | 6.7 kg/6.5kg* | Arduino MEGA + UNOs (encoders, pulse gen), TB6600 via NI USB-6001, MATLAB/LabVIEW |
| Symétrie SOLANO | 30/20 mm/s (X,Y/Z), 15/20 °/s (Rx,Ry/Rz) | ±18/±18/±6.5 mm / ±10°/±10°/±21° | Resolution 0.08–0.1 μm, repeatability ±0.25 μm | Height 104mm, Ø120 mm | 1 kg/5kg | ALPHA+ controller, Ethernet |
| PI H-811 (Miniature) | 20–25 mm/s(X,Y,Z) / ~28.6°/s(500 mrad/s) | ±17/±16/±6.5 mm / ±10°/±10°/±21° | MiM 0.08–0.2 μm / Repeatability ±0.06–0.15 μm | Height 114 mm, Ø136 mm | 2.2 kg/5kg | C-887 (EtherCAT/ TCP-IP/ RS-232) |
| Aerotech HEX150-125HL | 30 mm/s(X,Y) / 8 mm/s (Z) / 10°/s (Rx,Ry), 30°/s (Rz) | 42/44/17 mm / ±16° (Rx,Ry), ±42° (Rz) | MiM 15 nm / repeatability ±1.5 μm (X), ±0.4 μm (Z), ±3 arcsec | Height 125 mm,ϕ150 mm | 3 kg/5kg | HEX RC / Automation1 (TCP/IP ASCII, A3200 FireWire, HyperWire/ EtherCAT/ Modbus) |
| Newport / MKS HXP50 | 14/12/5 mm/s (X/Y/Z) / 6/6/15 °/s (Rx/Ry/Rz) | ±17/±15/±7 mm / ±9°/±8.5°/±18° | MiM 0.05–0.1 μm / repeatability 0.2 μm | Height 151 mm, Ø200 mm | 2.2kg/5 kg | HXP50-ELEC controller (TCP/IP) |
| PI (Physik Instrumente) H-820 | 20 mm/s (X,Y,Z) / 11.5°/s (200 mrad/s) | ±50/±50/±25 mm / ±15°/±15°/±30° | MiM 5 μm / repeatability ±1.5 μm (X,Y), ±0.5 μm (Z), ±8/±8/±25 μrad | Height 308 mm,Ø350 mm | 15 kg/20kg | C-887 6D controller(PI) |
| PI H-860(Linear-motor "Motion Simulator") | > 250 mm/s(X,Y,Z), 4 g,〜25 Hz(0.1°range) | ±7.5 mm / ±4° | — | Height 319 mm, Ø407 mm | 30 kg/1kg | PI 6D controller (TCP/IP, RS-232) |
| ACROME Stewart Platform | 10 mm/s (12 kg) / 40 mm/s (35 kg) / 12–40°/s (Roll/Pitch) | ±60〜100/±50〜103 mm / ±20〜30° | Resolution 100–200 μm / repeatability ±50–100 μm | Height 406–717 mm, Φ450mm | 14kg/12kg | TCP/UDP API (Ethernet, MATLAB/Simulink etc.) |
| Symétrie HEGOA | 200 mm/s(X,Y) / 120 mm/s(Z) / 50 °/s | ±100/±50 mm / ±23°/±23°/±30° | — | Height 420 mm, Ø500 mm | 30 kg/50kg | Ethernet(ALPHA+) |
| Mikrolar P1500 | 1,244 mm/s (X,Y) / 254 mm/s (Z) | Ø833 mm(XY)/ 315 mm(Z) / ±25°/±25°/±15° | Repeatability 25 μm | Height 833 mm, ϕ833 mm | 36 kg/150 kg | Software/UI(G-code) |

| | | | | | | |
|---|---|---|---|---|---|---|
| Quanser Hexapod | 0.67 m/s, 1 g, 0–10 Hz | ±7.4/±11/±5.4 cm / ±17°/±15°/±27° | Joint resolution 0.1 μm | Height 0.75 m, Φ=1.1m | 100 kg/100 kg | DAQ/amplifier,USB, QUARC (MATLAB/Simulink) |
| DOF Reality H6 (HERO 6) | 0.5 m/s / ~86°/s | ±17° | — | 1.20 × 1.50 × H 0.60 m | 90 kg | USB controll unit(SimRacingStudio) |

MiM: Minimum Incremental Motion
Load capacity depends on the stage height. The value shown corresponds to a stage height of 230 mm (the default configuration used in experiments). Increasing the stage height increases the load capacity.

**Supplementary Table 1.** Comparison of commercially available 6-DOF platforms. The maximum velocity of platform motion, range of motion, size, mass, load capacity, and control software or interface are compared.

##### 1.2. Mechanical design of 6AGM (CAD)

Here, we describe the mechanical design of the 6AGM system. The main components of the 6AGM include the stage, on which the animal is placed, the actuators (NEMA11 2PH, Pilang), and the connecting shafts linking the stage to the actuators. The stage, together with the connectors between the shafts and actuators, was designed in 3D CAD software (Fusion 360, Autodesk) and fabricated with a 3D printer (Form 2, Formlabs). Ball joints were implemented using magnetic sockets and steel balls, which were mounted with custom parts produced by 3D printing. Unlike conventional Stewart platforms, where actuators are vertically arranged and must bear much of the load, our design places linear actuators on a flat surface. This geometry allows the weight of the stage and its payload to be distributed primarily through normal forces rather than the actuators themselves, improving load support and reducing mechanical stress. Below, we present the final assembly along with each individual component and its description.

- 6dofPlatformAssembly.dxf

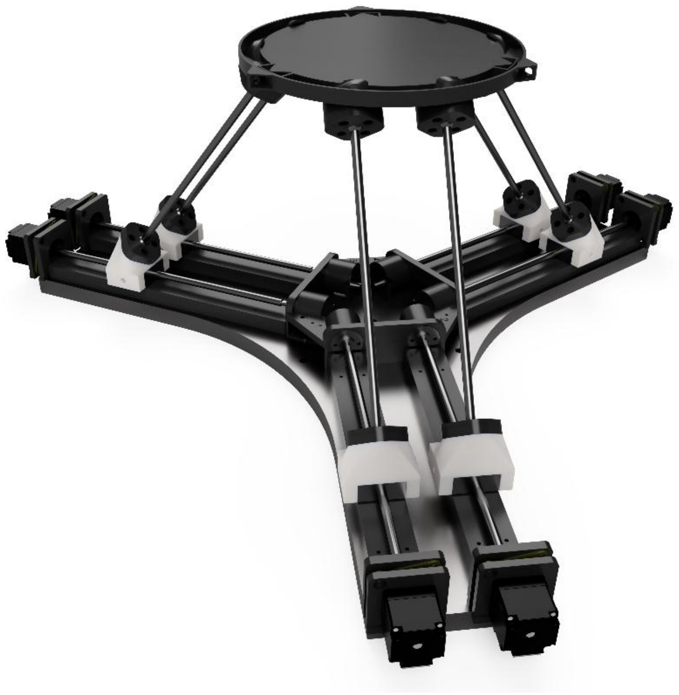

CAD model of the assembled 6AGM. For details of platform motion and operation.

➢ Stage.dxf

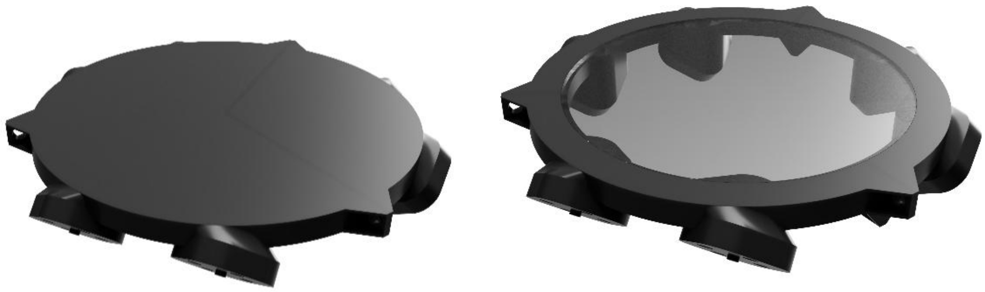

The stage, designed to accommodate head-fixed animals, is connected to six shafts via top joints. A circular opening, 14 mm in diameter, provides optical access for imaging the animal's underside with cameras. To reduce surface reflections from the acrylic plate, a black infrared-absorbing sheet (IR Flock Sheet; The Black Market) was applied when necessary.

➢ Base.dxf

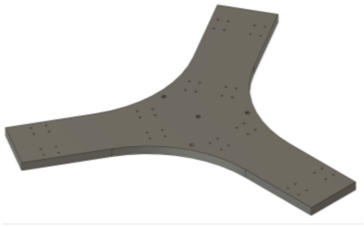

A 10-mm-thick aluminum plate was used to mount the six stepper motors. For precise stage control, the relative positions of the motors had to remain rigidly fixed, thereby ensuring stable and reproducible kinematics of the 6AGM.

➢ BallJointFixer.dxf

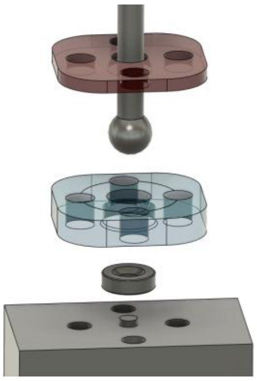

Ball joints connect each end of the shafts to the actuators and the stage. At the bottom, a magnetic socket allows the ball to roll smoothly, while the upper plate holds it in place without restricting motion. After 3D printing, the upper plate was trimmed to increase the range of motion. Oil (S4-T1500N, Sugiura Laboratory Inc.) was applied to all joints to minimize friction.

➢ FootJoint.dxf

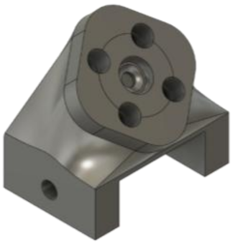

FootJoint connects the actuator and the BallJointFixer. The junction plane with the BallJointFixer is tilted to accommodate a higher range of motion.

➢ HeadPlateHolder.dxf

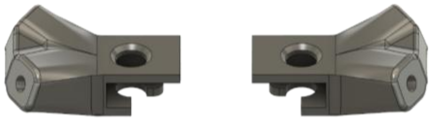

The head plate holder is fixed by rods attached at an angle to avoid interference with the platform below and the objective lens above.

➢ Photograph of the rotary encoder

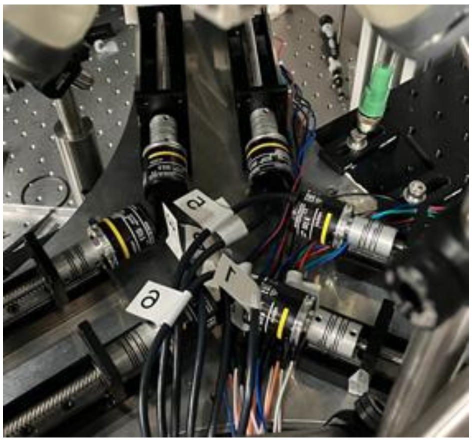

The inner ends of the linear guides are disassembled to connect rotary encoders (E6A2-CWZ3C) for tracking the rotations.

➢ Photograph of the damper.

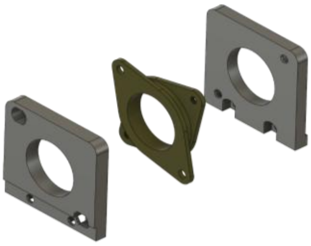

To prevent vibrations of the platform from propagating as severe artifacts during two-photon imaging, each stepper motor was mounted via a vibration-absorbing damper, attached with a custom adapter.

##### 1.3. Platform and stepper motor position coordinate conversion

This section outlines the coordinate transformation used to compute the positions of the six stepper motors from the platform's pose (position and orientation), thereby enabling user-specified platform motion. The actuators are controlled by solving the inverse kinematics associated with the desired stage trajectory. The base points of the ball joints $\boldsymbol{a_i}$ and the actuator directions $\boldsymbol{v_i}$ are defined in the equations below, expressed in the coordinate system of the base, with the origin at the center of the aluminum plate.

$$\boldsymbol{a_i} = \begin{bmatrix} \cos\theta_i & -\sin\theta_i & 0 \\ \sin\theta_i & \cos\theta_i & 0 \\ 0 & 0 & 1 \end{bmatrix} \begin{bmatrix} a_x \\ a_y \\ a_z \end{bmatrix} \quad \text{for } i = 2,4,6$$

$$\boldsymbol{a_i} = \begin{bmatrix} \cos\theta_i & -\sin\theta_i & 0 \\ \sin\theta_i & \cos\theta_i & 0 \\ 0 & 0 & 1 \end{bmatrix} \begin{bmatrix} a_x \\ -a_y \\ a_z \end{bmatrix} \quad \text{for } i = 1,3,5$$

$$\theta_i = 0,0,\frac{2\pi}{3},\frac{2\pi}{3},\frac{4\pi}{3},\frac{4\pi}{3}$$

$$\boldsymbol{v_i} = \begin{bmatrix} \cos\theta_i \\ \sin\theta_i \\ 0 \end{bmatrix} \quad \text{for } i = 1,2,\ldots,6$$

$$a_x = 80.5,\ a_y = 30.0,\ a_z = -25.5.$$

The ball joints connecting the stage and the shafts are represented in the following equations, expressed in the coordinate system of the stage, with the origin defined at the center of the stage surface.

$$\boldsymbol{b_i} = \begin{bmatrix} \cos\theta_i & -\sin\theta_i & 0 \\ \sin\theta_i & \cos\theta_i & 0 \\ 0 & 0 & 1 \end{bmatrix} \begin{bmatrix} b_x \\ b_y \\ b_z \end{bmatrix} \quad \text{for } i = 2,4,6$$

$$\boldsymbol{b_i} = \begin{bmatrix} \cos\theta_i & -\sin\theta_i & 0 \\ \sin\theta_i & \cos\theta_i & 0 \\ 0 & 0 & 1 \end{bmatrix} \begin{bmatrix} b_x \\ -b_y \\ b_z \end{bmatrix} \quad \text{for } i = 1,3,5$$

$$\theta_i = 0,0,\frac{2\pi}{3},\frac{2\pi}{3},\frac{4\pi}{3},\frac{4\pi}{3}$$

$$b_x = 197.2,\ b_y = 30.0,\ b_z = 47.7.$$

The actuator positions are calculated using the following equation.

$$l^2 = (\boldsymbol{R}\boldsymbol{b}_i + \boldsymbol{p}_i - \boldsymbol{a}_i - \lambda_i \boldsymbol{v}_i)^T (\boldsymbol{R}\boldsymbol{b}_i + \boldsymbol{p}_i - \boldsymbol{a}_i - \lambda_i \boldsymbol{v}_i)$$
$$l^2 = \lambda_i^2 - 2\boldsymbol{c}_i^T \boldsymbol{v}_i \lambda_i + \boldsymbol{c}_i^T \boldsymbol{c}_i$$
$$\lambda_i = \boldsymbol{c}_i^T \boldsymbol{v}_i + \sqrt{\left(\boldsymbol{c}_i^T \boldsymbol{v}_i\right)^2 - \boldsymbol{c}_i^T \boldsymbol{c}_i + l^2}$$
$$\text{where } \boldsymbol{c}_i = \boldsymbol{R}\boldsymbol{b}_i + \boldsymbol{p}_i - \boldsymbol{a}_i$$

where $\lambda_i$ denotes the position of actuator $i$, $\boldsymbol{R}$ and $\boldsymbol{p}$ represent the rotation matrix and the translation vector of the platform, respectively, and l is the length of the shafts.

To prevent parallel singularities, the six linear actuators were grouped into three adjacent parallel pairs, each aligned with one side of the triangular base. As initial open-loop control produced cumulative positioning drift, we implemented a high-speed feedback control to maintain accuracy over long experiments described in the next section.

##### 1.4. Discrete-time feedback–feedforward control scheme

To ensure accurate trajectory tracking of the 6-DOF platform, we implemented discrete-time feedback–feedforward control scheme (**Fig. 1e**). At each control step, the current positions of the six stepper motors were read out and compared with the ideal actuator positions, yielding offset values. These offsets were clipped to a tunable value to limit excessive commands and to prevent mechanical instability or oscillations. The offsets were subsequently added to the next feedforward increment $\Delta x_r = x_r[k+2] - x_r[k+1]$ calculated from the inverse kinematics of the planned trajectory, which is then converted to appropriate motor drive signals. By combining bounded feedback with predictive feedforward control, the platform was able to execute rapid and precise trajectories, including arbitrary tilts, rotations, and translations, while maintaining stability across all six degrees of freedom.

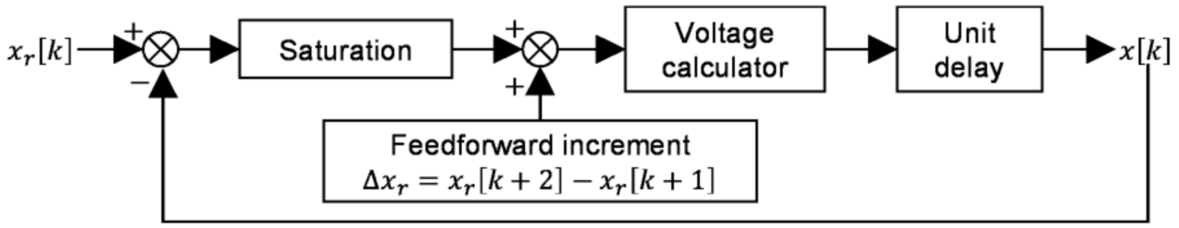


**Figure 1e (copy).**

##### 1.5. Control system of 6AGM

A schematic of the control system including real-time inverse kinematics and feedback control is shown in **Fig. 1c**.

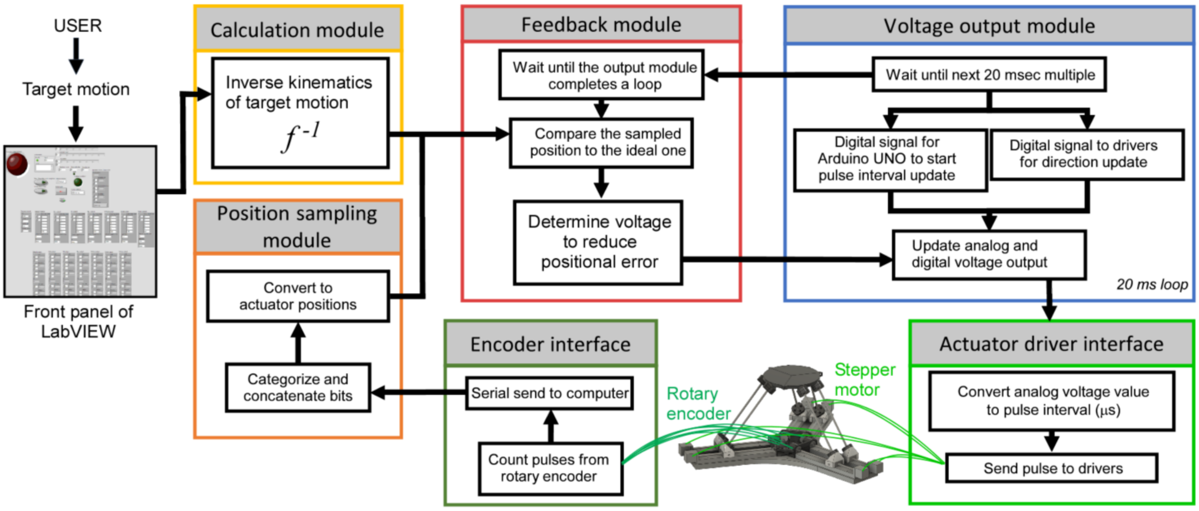


**Figure 1c (copy).**

The control system comprised four functional modules: the calculation module, position sampling module, feedback module, and voltage output module. The calculation module determined the analog voltage output required to execute user-specified platform trajectories, including transitions between motions with different initial configurations, and transmitted this information to the voltage output module. The position sampling module received serial transmissions from the encoder interface and converted these signals into actuator positions. The feedback module compared the estimated actuator positions with the reference values, computed updated outputs to compensate for deviations in subsequent iterations, and sent these corrections to the voltage output module.

Feedback control was implemented using rotary encoders (E6A2-CWZ3C) and an Arduino MEGA (16 MHz, 54 channels), which computed rotation angles and transmitted the data serially to the control PC at approximately 640 Hz. The voltage output module was designed to provide both analog and digital voltage outputs to the actuator driver interface (TB6600), as well as digital voltage outputs to the drivers via an NI USB-6001. The actuator driver interface consisted of six Arduino UNOs, which converted the analog signals into digital pulses to drive the stepper motors. The entire control program was developed in MATLAB (MathWorks) and LabVIEW (National Instruments).

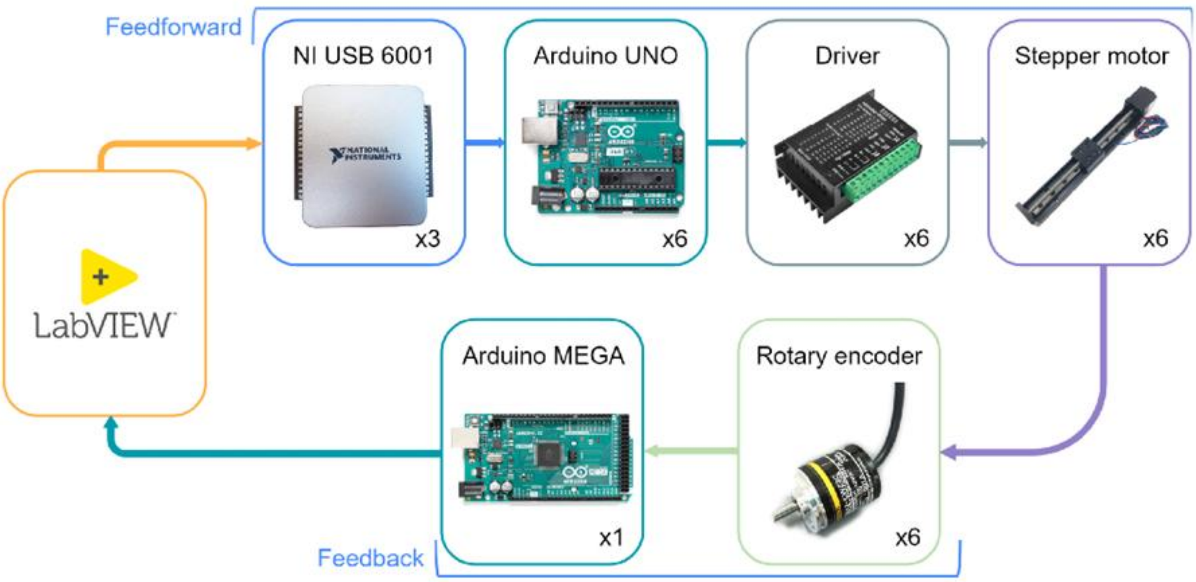


**Supplementary Figure 1 | Schematic of the control architecture for the 6AGM**

Schematic diagram of the control system architecture. Feedforward signals generated in LabVIEW were transmitted through NI USB-6001 devices (three units) to six Arduino UNOs, which provided digital control inputs to the actuator drivers (TB6600, six units). Each driver controlled a stepper motor (six units in total) for platform actuation. Feedback signals from six rotary encoders were collected by an Arduino MEGA, which transmitted the actuator position estimates to the control PC. This closed-loop configuration enabled precise trajectory execution through the combined feedforward and feedback pathways.

#### 1.6. Encoder interface

The Arduino MEGA board received three inputs from each rotary encoder: phase A, phase B, and phase Z. An interrupt was triggered on the rising edge of phase A, which incremented or decremented the pulse count depending on the phase offset of phase B. At higher rotation speeds, this operation could occasionally result in missed or inverted counts. To correct such errors, the encoder interface utilized the phase Z signal to realign the pulse count once per revolution. On the first detection of a Z pulse, the current pulse count was stored as a reference. For every subsequent Z activation, the counter was adjusted so that its difference from the reference corresponded exactly to an integer multiple of the encoder's pulses per revolution. This correction greatly improved stability over long durations, preventing drift entirely.

#### 1.7. Actuator driver interface

The actuator driver interface consisted of six Arduino UNO boards, each generating pulse signals for the corresponding motor drivers. The interface received a digital trigger from the control computer every 20 ms, initiating an analog voltage read. The measured voltage was then converted to a pulse interval (in microseconds) according to the following equation:

$$pulseInterval = round(\frac{B}{voltage+A}),$$

where

$$A = \frac{R \cdot t - r \cdot T}{T - t}, B = \frac{T \cdot t \cdot (R - r)}{T - t}, T = 10000, t = 20, R = 990, r = 0.$$

Here, $T$ and $t$ denote the upper and lower bounds of the pulse interval in microseconds, while $R$ and $r$ represent the bounds of the analog voltage reading. To correct for a consistent gain error between the Arduino UNO's ADC and the reference device (NI USB-6001), the upper bound of the analog voltage was set to 990 rather than the nominal 1023. Analog voltage values were returned with 10-bit resolution, whereas the fast PWM supported 16-bit resolution. This inverse mapping ensured that higher velocities were represented with finer precision. Importantly, under this mapping, constant noise in the voltage readout translates into a constant expected positional drift, independent of speed, thereby yielding more predictable long-duration behavior.

##### 1.8. Motion generation

By integrating the modules described above, we achieved smooth and stable motion of the platform. The static and dynamic parameters of the system were as follows. By default, the platform was positioned 30 mm below the head plate of the head-fixed mouse, with the stage center located beneath the occipital region (5 mm posterior to bregma). The platform was capable of rotations of up to 15° about the anterior–posterior axis (roll), 10° about the medio–lateral axis (pitch), and 15° about the vertical axis (yaw). Translational movements reached up to 165 mm horizontally and 115 mm vertically.

##### 1.9. LabVIEW software modules (Sub-VIs)

The LabVIEW program was divided into modules (Sub-VIs) to improve overall visibility and enable reuse of individual modules. The inputs, outputs, and functions of each Sub-VI are described below.

- ➢ Position_sampling.vi

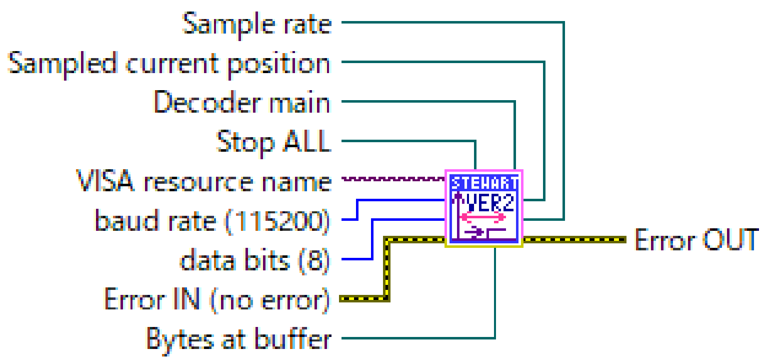


Reads data from serial port. This function decodes the received data to positions of each actuator. Once all the actuator positions are updated, the buffer at serial buffer is flushed to read the newest values.

➢ Calculate_motion.vi

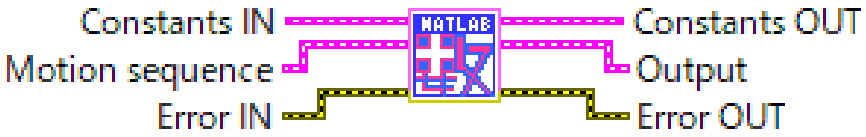


Calculates motion based on a time-series of six parameters. This function calculates the inverse kinematics of the platform and outputs a time-series of voltage values and actuator positions to be used by Output.vi and Feedback.vi, respectively.

➢ Initialization.vi

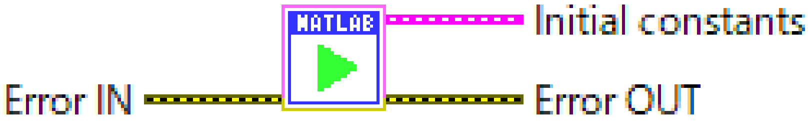


Initializes parameters used for inverse kinematics calculation

➢ Motion_TiltedRotation.vi

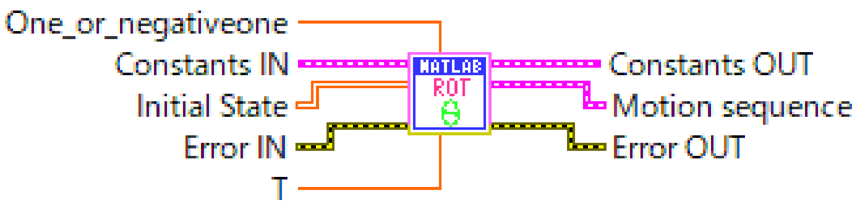


Generates a time-series of six parameters for tilted rotation motion.

➢ Motion_Roll.vi

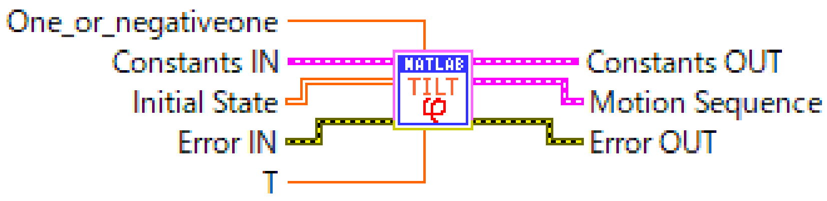


Generates a time-series of six parameters for roll motion.

➢ Motion_Yaw.vi

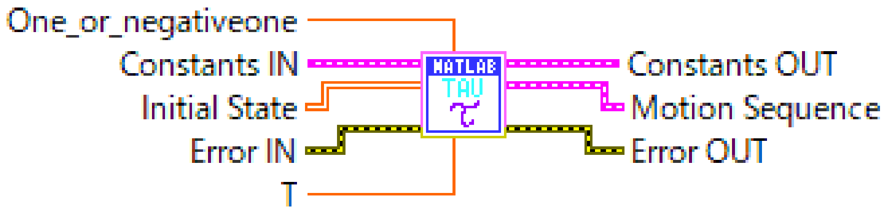


Generates a time-series of six parameters for yaw motion.

➢ Motion_Zvert.vi

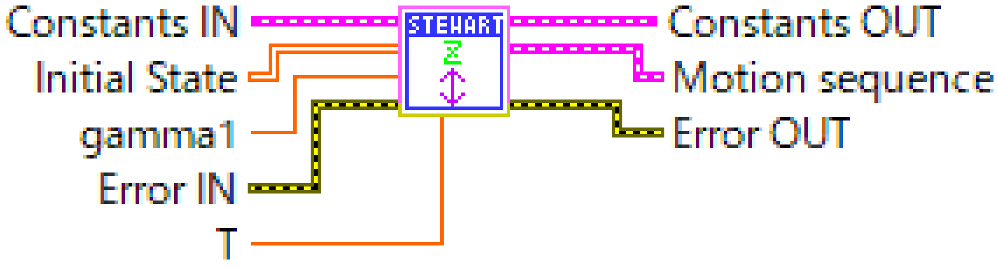


Generates a time-series of six parameters for up and down motion.

➢ Motion_CircularHor.vi

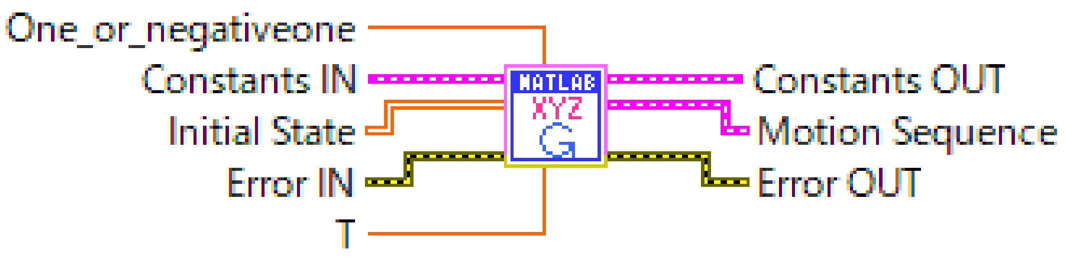


Generates a time-series of six parameters for horizontal circular motion.

➢ Motion_CircularVert.vi

Generates a time-series of six parameters for vertical circular motion.

➢ Output.vi

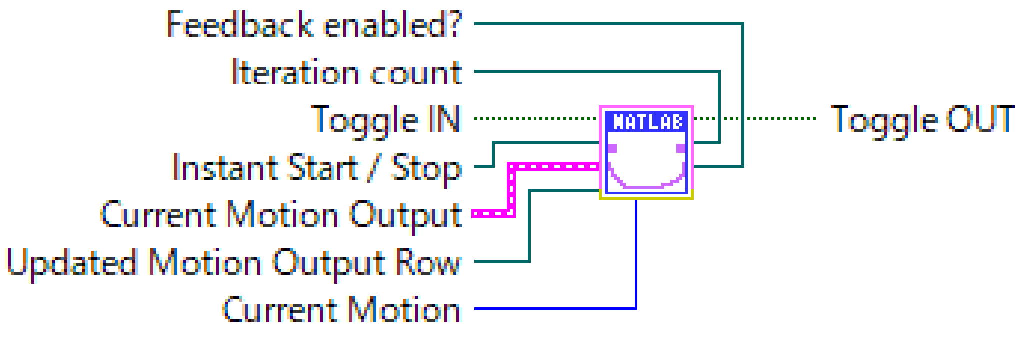


Iteratively updates analog and digital voltage applied from NI USB 6001. Every 20 ms, this vi updates the analog voltage applied to the actuator driver interface and the digital voltage applied to the motor drivers indicating the direction of motor rotation. Subsequently, it toggles its digital output to trigger an interrupt in the actuator driver interface that starts analog voltage read. This VI compares the current iteration value and the iteration value passed from the feedback module and selects the updated value if they match and default values from the calculation module otherwise.

➢ Feedback.vi

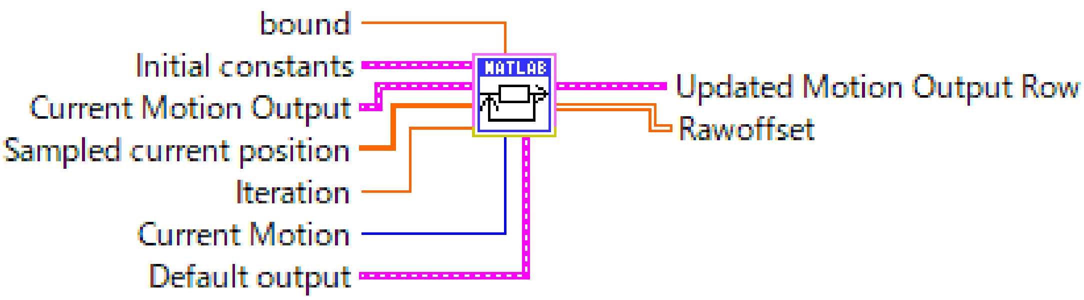


Compares the sampled position from Position_sampling.vi with the target trajectory from Calculate_motion.vi, and updates the voltage time series to compensate for the calculated offset. The feedback algorithm subtracts the observed actuator position from the reference position and clips it to a predetermined value. It then updates the feedforward increment by adding the clipped value to compensate for this offset in the next iteration output. The updated value goes through the voltage calculator and is used during the subsequent execution in Output.vi.

➢ Voltage_output_selector.vi

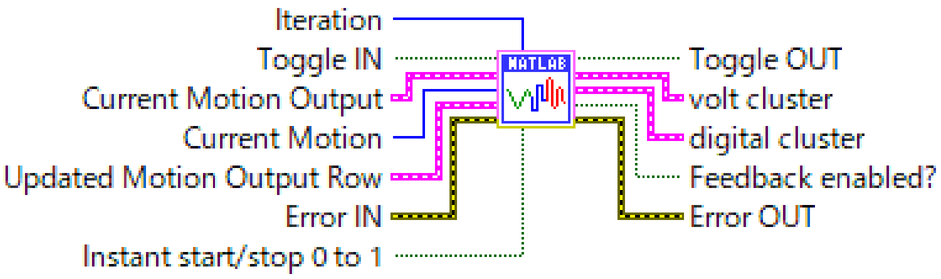


This VI resides inside Output.vi and selects the appropriate output.

If the iteration number tagged to the updated voltage from Feedback.vi matches the current iteration number of the Output.vi, this VI forwards the updated voltage value. Otherwise, this VI routes the voltage provided by Calculate_motion.vi. The selected voltage is sent out over NI USB-6001.

➢ Transition_motion.vi

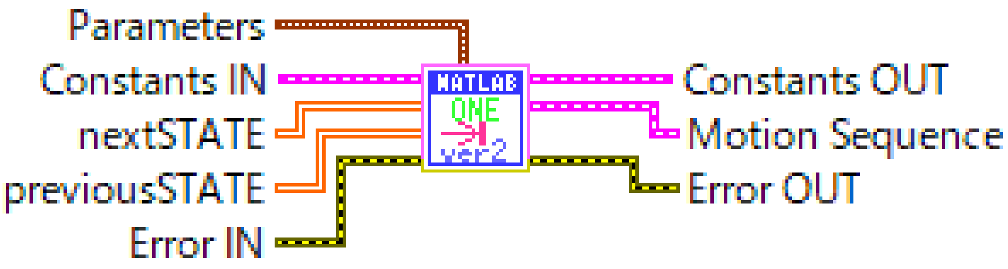


Generates time series of six parameters for transition motion. Transition motions are calculated by linearly interpolating the final state of the previous motion and the initial state of the next motion.

➢ ErrorHandler.vi

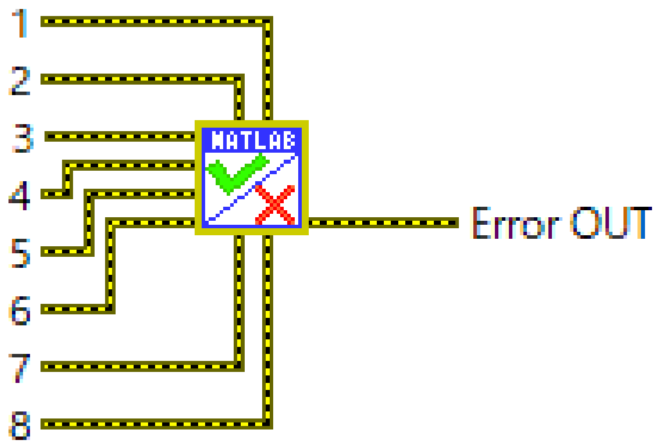


Handles errors and outputs a message indicating which VI threw an error.

#### 2. Camera calibration for 3D posture reconstruction

In our experiments, we observed the posture of an animal using four machine vision cameras. From these images, we identified the positions of multiple body parts using DLC; however, these positions are in two-dimensional camera sensor coordinates. Therefore, it is necessary to reconstruct the three-dimensional coordinates for each body part. In this section, we describe the procedure for calibrating the position of the camera sensor with respect to real-world coordinates using a checkerboard. By utilizing this calibration information, the positions of body parts identified by DeepLabCut can be accurately transformed into their corresponding locations in three-dimensional space.

**Workflow**

2.1. Single camera calibration

1. Checkerboard detection
2. Randomly select boards to use for parameter estimation
3. Obtain internal parameters assuming no radial distortion and project (normalize) coordinates
4. Obtain distortion parameters and undistort coordinates
5. Deproject undistorted coordinates
6. Obtain internal parameters
7. Assess fit performance of all boards
8. Repeat steps 3-7 using inlier boards obtained in step 7
9. Iterate steps 2-8 and save the model with best fit

2.2. Stereo camera calibration

1. Checkerboard detection
2. Obtain undistorted and projected coordinates for all cameras
3. Randomly select images to use for parameter estimation
4. Estimate the essential matrix
5. Calculate triangulation error of all boards
6. Repeat step 13-14 using inlier boards obtained in step 14
7. Estimate the essential matrix using inlier boards obtained in step 4-14
8. Repeat 12-16 and save the model with the lowest mean triangulation error
9. Repeat 12-17 for all camera pairs

###### 2.1.1. Checkerboard detection

We used a checkerboard (10 × 14, 19.25 mm × 19.25 mm) to capture calibration images. The checkerboard was filmed in a video at various angles. Around 1,000 images were uniformly sampled from the video to be clustered into 10 groups using k-means clustering. In total, 200 images were randomly selected from the clusters to be used for analysis.

###### 2.1.2. Randomly select boards to use for parameter estimation

We randomly select a fixed number of checkerboards out of 200 images to be used for parameter estimation.

Four cameras were positioned around the platform along with 4 infrared light sources. The platform was covered by an infrared absorptive cloth (IR Flock Sheet, The black market) to minimize reflection.

###### 2.1.3. Intrinsic and extrinsic parameter estimation

$$\hat{q} = \frac{1}{Q_z}\begin{bmatrix}\alpha_x & s & q_{0x} & 0\\ 0 & \alpha_y & q_{0y} & 0\\ 0 & 0 & 1 & 0\end{bmatrix}\begin{bmatrix}R & p\\ 0 & 1\end{bmatrix}\begin{bmatrix}Q^\circ\\ 1\end{bmatrix}$$

$$= \frac{1}{Q_z}KQ$$

where

$$K = \begin{bmatrix}\alpha_x & s & q_{0x}\\ 0 & \alpha_y & q_{0y}\\ 0 & 0 & 1\end{bmatrix}$$

$$Q = \begin{bmatrix}R & p\\ 0 & 1\end{bmatrix}\hat{Q}^\circ$$

Zhang's method (Z. Zhang 2000) was used for intrinsic and extrinsic parameter estimation of pinhole camera model. In this step, we naively estimated the 6 parameters of camera matrix K assuming no radial distortion in order to obtain projection of the checkerboard vertices onto a normalized image plane.

###### 2.1.4. Obtain distortion parameters and undistort coordinates

$$\bar{q_i} = q_0 + \frac{q_i - q_0}{1 + k_1\|q_i - q_0\|^2 + k_2\|q_i - q_0\|^4}$$

$\bar{q_i}$ is the ideal distortion free coordinates (projected to $z = 1$)

$q_i$ is the real distorted coordinates (projected to $z = 1$)

$q_0$ is the distortion center (not necessarily equal to the principal point)

The division model with two parameters was employed to model radial distortion of cameras. This model was used since it requires fewer terms than the polynomial model used in Zhang's method to model moderate to severe distortions.

The two-parameter division model assumes that the camera lens distorts a straight line on the projected plane to a circular arc. Therefore, the optimization of the parameters involves fitting an arc to distorted colinear points. Contrary to Zhang's iterative method that tries to minimize distance between projected points and distortion free points, this model uses the relative positions of the projected points, making it more robust to fluctuations in projection accuracy.

In this step, we first optimize radial distortion using one parameter division model to obtain k1 and q0. Subsequently, we use two-parameter division model with a fixed value of q0 from one-parameter estimation and obtain k1 and k2.

**2.1.5 Deprojection of undistorted coordinates**

After distortion parameter estimation, the projected coordinates of the checkerboards are undistorted and deprojected back to pixel coordinates.

**2.1.6 Obtain internal parameters**

Similarly to step 4-3, we obtain internal parameters assuming the distortion is corrected.

**2.1.7 Assess fit performance of all boards**

Using the parameters obtained in steps 2.1.4 through 2.1.6, we undistort and project all the boards and calculate the distance between the projected points and ideal points.

**2.1.8 Repeat steps 2.1.3-7 using inlier boards obtained in step 2.1.7**

We use all checkerboards with mean projection error lower than threshold value to calculate camera parameters. We then calculate the mean projection error of these inlier boards.

**2.1.9 Iterate steps 2.1.2-8 and save the model with best fit**

We iterate steps 2-8 and save the model with the lowest mean projection error of inlier boards.

**2.2.1 Checkerboard detection**

We used a checkerboard (2 × 3, 16 mm × 16 mm) to capture calibration images. Similarly to step 4-1, the checkerboard was filmed in a video at various angles. Around 2000 images were uniformly sampled from the video to be clustered into 10 groups using k-means clustering. In total, 1000 images were randomly selected from the clusters to be used for analysis. Around 200 images were used for each camera pair.

**2.2.2 Obtain undistorted and projected coordinates for all cameras**

Using the parameters obtained in step 2.1.4-9, undistort and project all the images onto the normalized image plane.

**2.2.3 Randomly select images to use for parameter estimation**

Randomly select 10 boards for reconstructing the essential matrix for a camera pair.

**2.2.4 Estimate the essential matrix**

We estimate the essential matrix of the chosen camera pair using a normalized 8 point algorithm.

**2.2.5 Calculate triangulation error of all boards**

Using the parameters obtained in 2.1.4-2.2.4, 3D coordinates along with their triangulation error are calculated for all the boards.

**2.1.6 Repeat step 2.2.4-5 using inlier boards obtained in step 2.1.5**

Using all the boards with triangulation error lower than threshold value, the essential matrix is recalculated.

**2.1.7. Calculate triangulation error of all inlier boards**

The recalculated essential matrix is used to obtain the triangulation error of the inlier boards.

The mean triangulation error is subsequently calculated.

**2.1.8. Repeat 2.2.3-7 and save the model with the lowest mean triangulation error**

We iterate steps 2.2.3-7 and save the model with the lowest mean triangulation error of inlier boards.

##### 2.1.9 Repeat 2.2.3-9 for all camera pairs

Steps 2.2.3-8 are repeated for all the camera pairs.

After estimating camera parameters, the camera pairs underwent stereo calibration to estimate their relative positions. Normalized 8-point algorithm was used for estimation. In order to filter out inaccurately detected checkerboards, RANSAC algorithm (Fischler and Bolles 1981) was employed.
